# ANCHOR: haplotype-aware allelic and isoform inference from single-cell long-read RNA sequencing with de novo variant calling

**DOI:** 10.64898/2026.06.08.730656

**Authors:** Zhi-Can Fu, Chuanxin Zhang, Yelian Yan, Yuxin Xu, Xiaoyu Yin, Tietuo Tao, Peng Lu, Yalin Liang, Haiyang Wu, Wentao Cui, Runtong Hou, Xuepeng Chen, Yuwen Ke, Yanqiang Li, Zi-Jiang Chen, Tao Huang, Keliang Wu, Shenli Yuan

## Abstract

Long-read RNA sequencing enables haplotype– and isoform-resolved allelic analysis of transcriptomes, yet extending this capability to single cells and distinct cell types remains computationally challenging due to sparse coverage, sequencing errors, incomplete variant information, and reference-biased transcript assignment. Here we present ANCHOR, a haplotype-aware framework for single-cell long-read RNA sequencing that performs de novo expressed-variant discovery, molecule-level haplotype assignment and isoform-resolved allelic quantification. ANCHOR combines a signed-graph variant caller, pair hidden Markov modelling and beta-binomial UMI aggregation to infer parental allele counts for genes and splice-resolved isoforms, without requiring a pre-existing phased genotype or deep learning. In human single-cell long-read RNA benchmarks, ANCHOR improved variant-calling performance over tested long-read RNA callers at single-cell and low-to-moderate coverage, and its beta-binomial model reduced depth-driven false positives in allele-specific expression testing. Applied to newly generated single-cell long-read RNA-seq data from reciprocal mouse crosses during gastrulation, ANCHOR resolved cell-type– and isoform-specific parent-of-origin imprinting and identified an antagonistic maternally biased *Sgce* isoform. ANCHOR provides a general framework for allele-and isoform-resolved analysis of diploid single-cell long-read transcriptomes.

## Introduction

Allele-specific expression (ASE), in which the two parental chromosomes contribute unequally to the transcriptome, provides a readout of cis-regulatory variation, X-chromosome inactivation and genomic imprinting^1–4^. In single-cell short-read data, however, ASE is usually inferred from isolated SNPs covered by few reads. These local measurements cannot be linked back to the full-length RNA molecule or its splice structure, making allele-specific isoform usage difficult to resolve^5–8^. Single-cell long-read sequencing can, in principle, address both limitations^9,10^. Because an Oxford Nanopore (ONT)^11^ or PacBio^12^ read may span a full-length cDNA molecule, it can sample several heterozygous sites from the same transcript and connect them to both cell identity and splice structure. However, this capability depends on three unresolved computational steps. Informative heterozygous sites are often not known in advance, making many analyses dependent on a matched or pre-existing genotype. ONT base-calling errors and limited read support per unique molecular identifier (UMI) make reference/alternate counting noisy, while high read depth can make small technical skews statistically significant despite limited biological effect size. Finally, assigning reads to isoforms against a single reference transcriptome can misclassify molecules from the non-reference allele when the two alleles differ in splice structure, thereby obscuring the allele-specific isoform events that the analysis is designed to resolve (Methods, “Isoform assignment”).

Existing long-read methods address related but incomplete aspects of allele– and isoform-resolved transcriptome analysis. LongAllele^13^ uses joint variant–haplotype–read EM to infer allelic expression and transcript usage, although a runnable implementation was not available for direct comparison at the time of writing. LongcallR^14^ performs variant calling, phasing and ASE analysis in bulk long-read RNA-seq. LORALS^15^ and isoLASER^16^ quantify isoform-level ASE in human tissues, but rely on paired DNA genotypes for phasing. To our knowledge, no existing single-cell long-read workflow jointly supports RNA-only heterozygous SNP discovery, molecule-level haplotype assignment, isoform-resolved ASE and reciprocal-cross parent-of-origin decomposition. We present ANCHOR (Allelic iNference in single-Cell Haplotype-aware lOng-Reads), an interpretable framework for haplotype-aware allelic and isoform inference from single-cell long-read RNA-seq. ANCHOR discovers expressed heterozygous variants de novo with a signed-graph caller, assigns molecules to parental haplotypes using a pair hidden Markov model (pair-HMM), and aggregates read evidence within UMIs through a Beta-binomial model that can abstain when allelic evidence is ambiguous. Isoforms are defined from observed splice structure rather than reference transcript assignment, allowing allele-specific splice patterns to be retained. The method requires a diploid sample with sufficient expressed heterozygosity, mappability and long-read coverage, but not a pre-existing phased genotype.

We applied ANCHOR to newly generated single-cell long-read RNA-seq data from reciprocal F1 hybrid mouse embryos across gastrulation (E6.5–E8.5). Reciprocal crosses distinguish parent-of-origin effects from strain-biased cis regulation because true imprinting, but not strain bias, inverts with cross direction^17,18^. By applying this logic at single-cell and isoform resolution, ANCHOR resolved cell-type– and isoform-specific imprinting, including a maternally biased transposon-derived *Sgce* isoform that opposes the canonical paternal *Sgce* isoform.

## Results

### ANCHOR infers allele– and isoform-resolved expression from single-cell long reads

ANCHOR takes a position-sorted single-cell long-read BAM and produces per-cell, per-gene and per-isoform parental allele counts (**Fig. 1a**). Reads are piled into a molecule-by-site sign matrix. ANCHOR first discovers heterozygous sites de novo from this matrix with a two-stage gradient-boosted classifier (below), then, for each read, a pair-HMM scores the read against the two haplotype reference sequences by global forward dynamic programming, yielding a calibrated per-site log-likelihood ratio (LLR). A read-level HMM integrates LLRs along the molecule, a Beta-binomial model aggregates reads within each UMI into a four-state posterior (allele 1, allele 2, mixed, unknown), and UMIs are summed per cell and per isoform. Isoforms are defined by observed splice junctions, not by reference mapping, so allele-specific splicing is preserved rather than collapsed onto the reference transcript (Supplementary Note 1). In practice, ANCHOR first decides which sites are informative, then which haplotype each molecule supports, then aggregates those molecule-level decisions into cell– and isoform-level allelic counts. The framework is organism– and dataset-agnostic, learning haplotype identity de novo so that no pre-existing genotype is required.

**Fig. 1.**
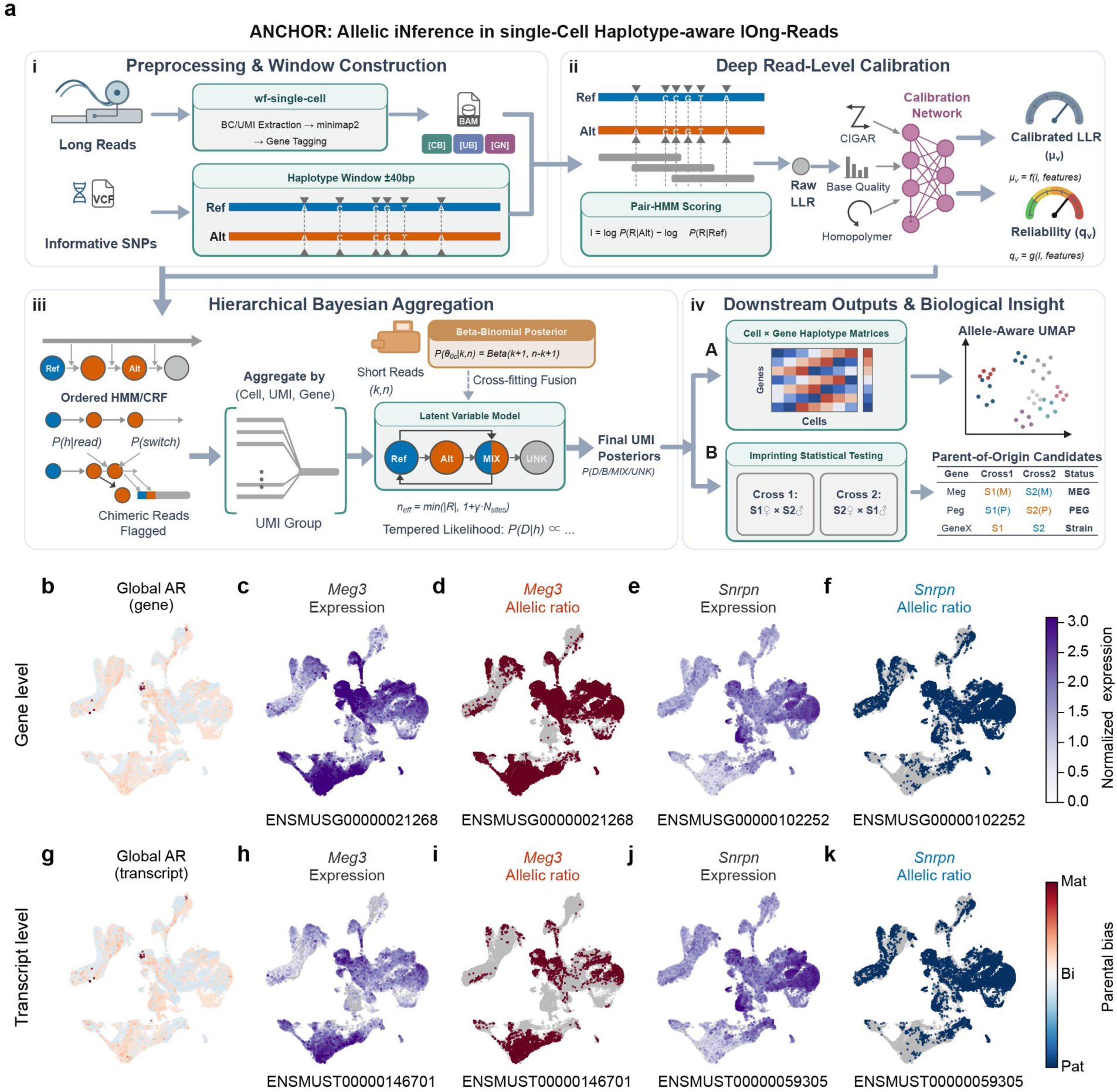
| ANCHOR resolves single-cell, haplotype– and isoform-resolved allelic expression. (**a**) The ANCHOR pipeline: per-site pair-HMM emission likelihoods, batch-aware Platt calibration, and a per-read hidden Markov model followed by UMI-level Beta-binomial aggregation and cell-level summation that outputs allele-1 and allele-2 counts per cell, gene and isoform. (**b**–**f**) Gene-level and (**g**–**k**) transcript-level measures projected onto a Harmony-integrated UMAP of 55,729 single cells profiled by long-read RNA sequencing from hybrid mouse gastrulating embryos. (**b**,**g**) Per-cell global allelic ratio. (**c**,**e**,**h**,**j**) Log-normalized expression of *Meg3* and *Snrpn*; grey, zero raw UMIs. (**d**,**f**,**i**,**k**) Per-gene and per-isoform parental-bias score = (maternal − paternal)/(maternal + paternal), where the maternal allele is assigned according to each cross direction; grey, fewer than four allele-informative UMIs (a pre-specified visualization filter). Colour scales: expression, Purples; parental bias, blue (paternal) to white (biallelic) to red (maternal); *n*, number of coloured cells, is shown per panel. *The two example genes shown (Meg3, Snrpn) illustrate the gene– and isoform-level expression and parental-bias output and are not presented as a biological result (see Results)*.

As an illustration of the pipeline output, ANCHOR’s per-cell, per-gene and per-isoform allelic calls project onto a single-cell embedding (Fig. 1b–k). The genome-wide allelic ratio is near-balanced, with no systematic mapping bias (Fig. 1b,g). Together, these panels illustrate ANCHOR’s end-to-end output: a near-balanced, genotype-free allelic readout resolved simultaneously at cell, gene and isoform level.

### ANCHOR’s de novo variant caller outperforms existing callers on human HG002

To discover heterozygous variants without a pre-existing genotype, the ANCHOR caller works from the reads alone. Viewing the molecule-by-site sign matrix *A* ∈ {+1, −1, NaN} as a signed bipartite graph, a robust rank-1 two-colouring *A* ≈ *hδ*^T^ + *R* assigns each molecule a haplotype sign *h* and each site a phase sign δ. The residual sign conflict (sites that resist this two-haplotype split) together with graph-connectivity features feed a two-stage gradient-boosted classifier that emits a phased VCF (**Fig. 2a**). The factorization is well behaved at scale. On a chromosome-11 pilot of 4.15 million molecules by 1.34 million sites (2.44 billion observed entries), the alternating optimization converges within eight iterations to 99.61% weighted agreement (0.41% residual; Extended Data Fig. 1a), with per-site frustration cleanly separating true variants from artifacts (Extended Data Fig. 1b,c) and graph-negative mass showing the same separation un-normalized (Extended Data Fig. 1d), and the two-colouring reproduces the observed sign structure with 99.5% agreement on real data (Fig. 2b), showing that the rank-1 factorization recovers haplotype phase directly from the sign matrix at chromosome scale.

**Fig. 2.**
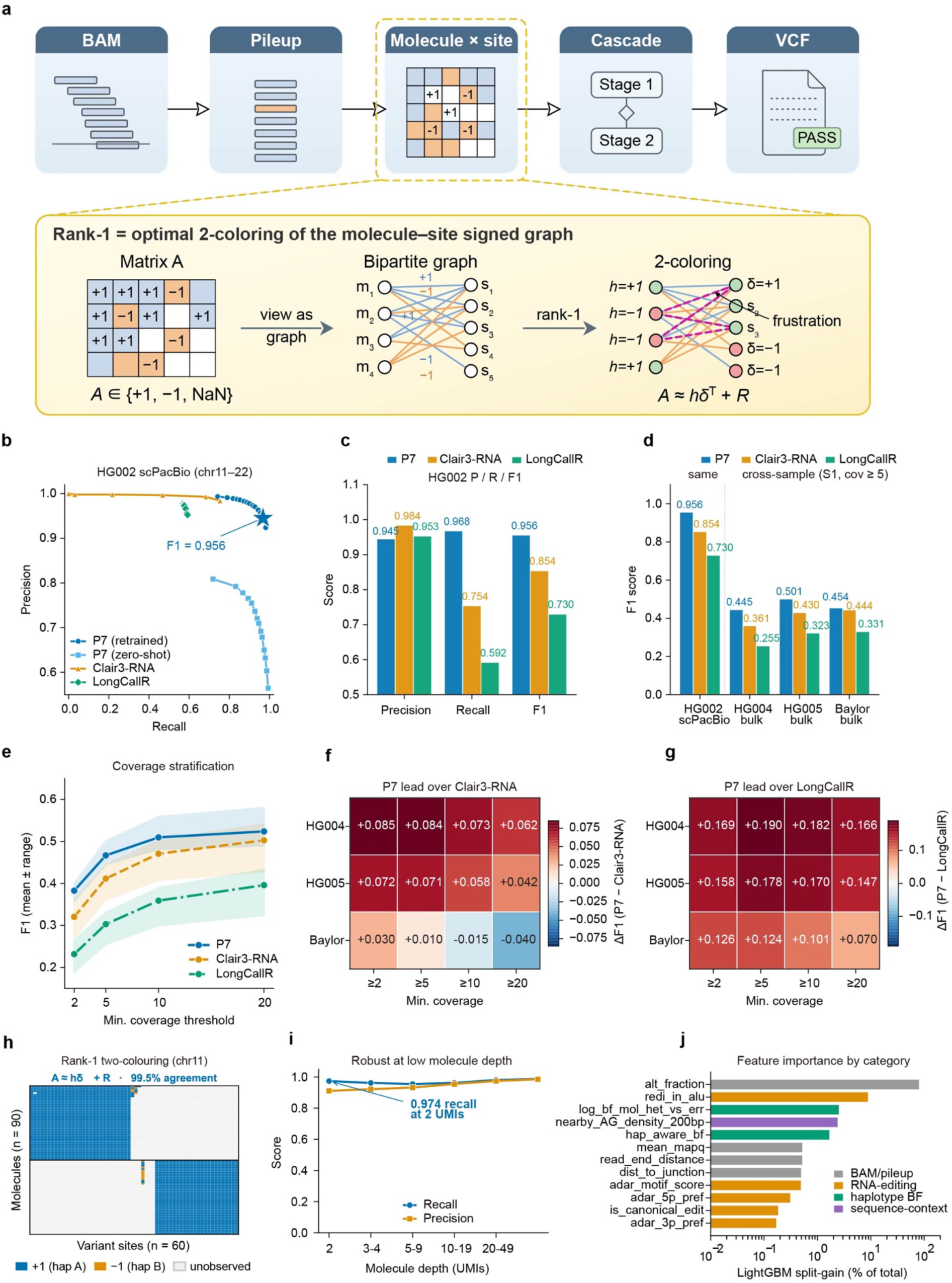
| De novo single-cell long-read variant calling and its benchmarking on human HG002. (**a**) The ANCHOR caller: the molecule-by-site sign matrix *A* ∈ {+1, −1, NaN} (one entry per molecule × site) is factorized into a molecule-haplotype vector *h*, a site-phase vector *δ* and a residual *R* (*A* ≈ *hδ*^T^ + *R*, a rank-1 two-colouring); the sign of each *h* entry assigns its molecule to one of two haplotypes, after which a two-stage classifier emits a phased VCF. (**b**) A representative chromosome-11 observed molecule-by-site sub-matrix reordered by *h* and *δ*, annotated with the fraction of entries whose sign matches the haplotype/phase assignment from **a**. (**c**) Precision–recall on the HG002 single-cell PacBio held-out split (chr11–22) for ANCHOR (trained chr1–10), ANCHOR with mouse-trained weights applied zero-shot, Clair3-RNA and LongCallR; the star marks best-F1. (**d**) Precision, recall and F1 at best-F1, by method (ANCHOR 0.945/0.968/0.956; Clair3-RNA 0.984/0.754/0.854; LongCallR 0.953/0.592/0.730). (**e**) Recall and precision versus molecule (UMI) depth on the HG002 split (recall ≥ 0.95 across depths; 0.974 at two-UMI sites). (**f**) F1 across four samples — HG002 single-cell PacBio (same-sample) and three cross-sample bulk Iso-Seq datasets (HG004, HG005, Baylor HG002) at minimum coverage ≥ 5. (**g**) Mean F1 across the three cross-sample datasets versus minimum coverage; band, per-sample range. (**h**,**i**) F1 advantage of ANCHOR over Clair3-RNA (**h**) and LongCallR (**i**) per sample and coverage. (**j**) Caller feature importance by evidence category.

On human HG002 single-cell PacBio data, a cross-species, cross-platform test against the GIAB truth set^19^, ANCHOR reaches precision 0.945, recall 0.968 and F1 = 0.956 on a held-out chromosome split (chr11–22, trained on chr1–10; all evaluated methods scored on the same candidate set, Methods), exceeding Clair3-RNA (F1 = 0.854) and LongCallR (F1 = 0.730), driven by higher recall at matched precision (Fig. 2c,d). Within the chr1–10 training set, performance is stable across five-fold cross-validation (validation-fold AUPRC 0.847 ± 0.002; Extended Data Fig. 2a–d), and it is uniform across held-out chromosomes (mean F1 = 0.957, range 0.932–0.971). Recall stays above 0.95 even for two-UMI sites (0.974), the lowest depth at which a heterozygous site can be called (Fig. 2e). Model-variant ablation on HG002 confirms the production configuration as the highest-F1 setting (Extended Data Fig. 2e). Across three independent bulk Iso-Seq datasets^20^ (HG004, HG005, Baylor HG002), used as a cross-individual generalization test, ANCHOR leads on coverage-averaged F1 across the full coverage range (Fig. 2f,g; Extended Data Fig. 2f). The only exception is the bulk Baylor sample at coverage ≥ 10, where Clair3-RNA overtakes ANCHOR as its cell-level features lose signal in bulk data (Fig. 2h; Extended Data Fig. 2g,h). On the single-cell data, where those cell-level features apply, ANCHOR leads at all coverages. Feature attributions show the caller weighs graph-linkage, sequence-context and BAM-quality evidence in balance (**Fig. 2j**). Together, these single-cell and cross-individual bulk tests show that ANCHOR’s de novo caller transfers to human single-cell PacBio data and generalizes across individuals on bulk Iso-Seq, exceeding Clair3-RNA and LongCallR on F1 while remaining accurate down to two-UMI sites.

### ANCHOR’s caller and aggregation engine transfer to a mouse reciprocal-cross cohort

To test ANCHOR on a reciprocal-cross design where parent-of-origin and strain effects can be separated, we profiled reciprocal C57BL/6J × DBA/2 crosses by sibling multi-omics. Quality control confirms high long-read sequencing quality, the marker panel and a coverage advantage over Illumina (Extended Data Fig. 3a–j), and the cells integrate into a joint single-cell atlas with matched per-platform QC (Extended Data Fig. 4a–h). Reciprocal direction inversion supplies the biological decision rule used throughout: a true parent-of-origin effect flips sign between the forward and reverse crosses, whereas a cis-strain effect keeps the same B6/DBA sign in both. On the in-house mouse reciprocal-cross scONT data, an ablation across operating points shows that each model layer adds signal (AUPRC 0.909, 0.939, 0.970 and 0.976 for the BAM-only, +context, +cascade and +graph layers, respectively). The calibrated cascade is the primary driver, and the graph/linkage layer adds recall specifically at the high-precision operating point (recall rising from 0.29 to 0.37 at s2 ≥ 0.95), where de-novo calling is otherwise conservative (Fig. 3a; Extended Data Fig. 1e–g). The same caller leads a six-method cross-sample comparison (its four cumulative layers, Clair3-RNA and LongCallR; Fig. 3b) and is accurate across three forward and three reverse crosses (Fig. 3c), with stage-1 attributions dominated by graph-derived features (Fig. 3d), precision and recall uniform per chromosome across the 18 autosomes (chr11 held out; Fig. 3e) and graph, context and BAM-quality evidence forming the top-ranked features (Fig. 3f; Extended Data Fig. 1h). The caller is also robust to RNA-editing, which must be distinguished from true DNA variants in RNA-based variant calling^21^, and other artifact contexts (Extended Data Fig. 5a–e), maintains recall across variant allele fraction, BAM coverage and supporting-cell counts (Extended Data Fig. 5f–h), and is well calibrated at both the final and gate scores (Extended Data Fig. 5i,j). Together, the same de novo caller validated on human HG002 leads a six-method comparison on mouse scONT, is uniform across all 18 autosomes, and stays robust to RNA-editing and other artifact contexts.

**Fig. 3.**
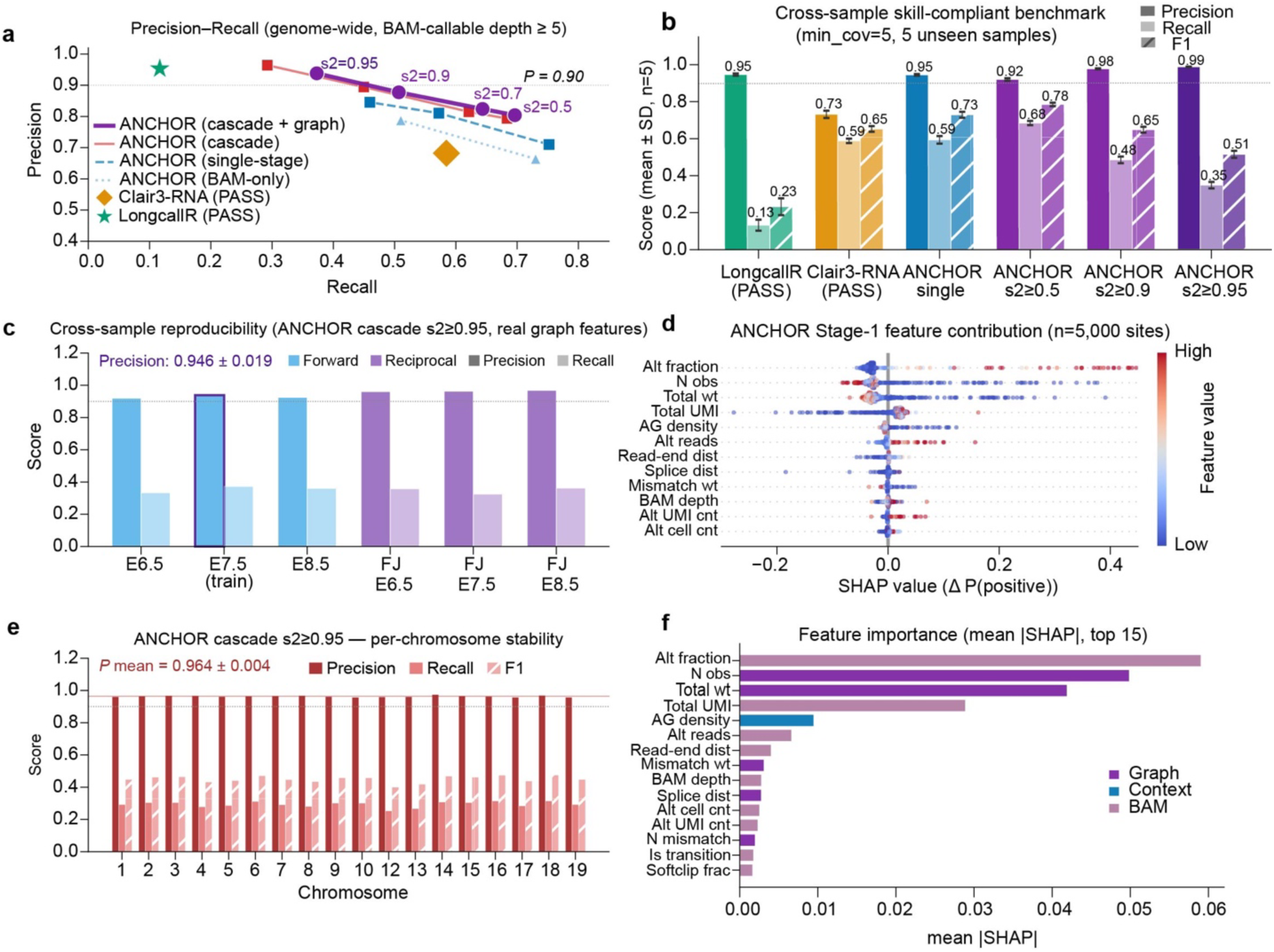
| ANCHOR de novo variant calling on the in-house mouse reciprocal-cross scONT dataset. (**a**) Precision–recall on chr1–chr19 (chr11 held out) at BAM-callable depth ≥ 5; ANCHOR variants are score-threshold sweeps for four cumulative model layers (BAM-only, single-stage, the two-stage cascade, and cascade plus graph/linkage features); Clair3-RNA and LongCallR are single PASS points. (**b**) Cross-sample precision, recall and F1 for six methods at minimum coverage ≥ 5 across five held-out samples (mean ± s.d.; the sixth sample was used for training and is excluded here, cf. **c**); precision is depth-corrected. (**c**) Per-sample precision and recall for ANCHOR (cascade + graph) at s2 ≥ 0.95 across three forward and three reciprocal crosses; the training sample carries a dark outline. (**d**) SHAP beeswarm for the stage-1 model (*n* = 5,000 sites). (**e**) Per-chromosome precision, recall and F1 for ANCHOR (cascade, no graph) at s2 ≥ 0.95 across 18 autosomes (chr11 held out). (**f**) Top-15 features by mean |SHAP|, coloured by family (graph, context, BAM).

With informative sites called, we next validated the second ANCHOR module: the molecule-level aggregation engine that turns calibrated per-read evidence into cell– and isoform-level counts. Hierarchical aggregation measurably improves allelic recovery over read pooling. On semi-synthetic molecules carrying known per-UMI labels (270 synthetic BAMs built from real scONT reads; Extended Data Fig. 6a), mean UMI-label recovery is 99.58% ± 0.30% across SNP-per-UMI and read-per-UMI grids (Extended Data Fig. 6b,c), insensitive to allelic ratio (0.50 vs 0.99; Extended Data Fig. 6d) and to Beta-binomial overdispersion (Extended Data Fig. 6e). Against matched Illumina, UMI-aware aggregation reduces allelic-calling error ∼47-fold in the diagnostic two-UMI regime relative to a read-level majority-vote ablation that ignores UMI structure, with smaller fold-reductions at higher UMI counts (Extended Data Fig. 7a). The pair-HMM emission model makes 41.4 million UMIs decodable in E6.5, versus 14.4 million for a genome-wide SNP-counting baseline (2.88×; Extended Data Fig. 7d), because it can score reads in SNP-sparse regions that counting cannot. Pseudo-bulk gene-level allelic ratios agree closely with matched Illumina (Extended Data Fig. 7e). Abstention is well-targeted: the Beta-binomial posterior accepts 98.9% of UMIs at 99.5% accuracy, while the 1.1% it sets aside would be only 81.5% accurate if forced to call. This confirms that abstention isolates genuinely ambiguous molecules (Extended Data Fig. 7c). Together, these results show that ANCHOR’s de novo caller and aggregation engine both transfer to the mouse reciprocal-cross cohort, with the caller ranking first in the six-method comparison and the engine recovering allelic state accurately at low molecule counts.

### ANCHOR controls depth-driven false positives for robust-effect ASE

To test whether ANCHOR separates genuine allelic imbalance from sampling noise, we benchmarked ANCHOR against scDALI-Hom^22^, ASPEN^23^ and MBASED^24^ on an independent truth set defined entirely from the matched Illumina data, with imprinted genes identified by reciprocal-cross direction inversion (3,676 genes: 186 imprinted, 761 biallelic-null, 2,729 strain-biased; **Fig. 4a**). On the biallelic-null negative controls (genes with no reproducible allelic skew at a matched-Illumina allelic ratio of 0.35–0.65 in both crosses), ANCHOR called 2 of 759 testable genes significant at FDR < 0.05, whereas scDALI-Hom, ASPEN and MBASED called 256/739, 223/473 and 272/717 on their respective testable sets (Fig. 4b; Extended Data Fig. 8a). Imposing a matched effect-size floor (|AR − 0.5| ≥ 0.10) on the competitors collapses their false-positive rate to 5–6% (versus ANCHOR’s 0.26%, 2 of 759) while leaving their imprinted recall at 58–68%, comparable to ANCHOR’s 57.5% (Extended Data Fig. 8b–d). ANCHOR’s contribution is reaching this specificity without a tunable effect-size threshold.

**Fig. 4.**
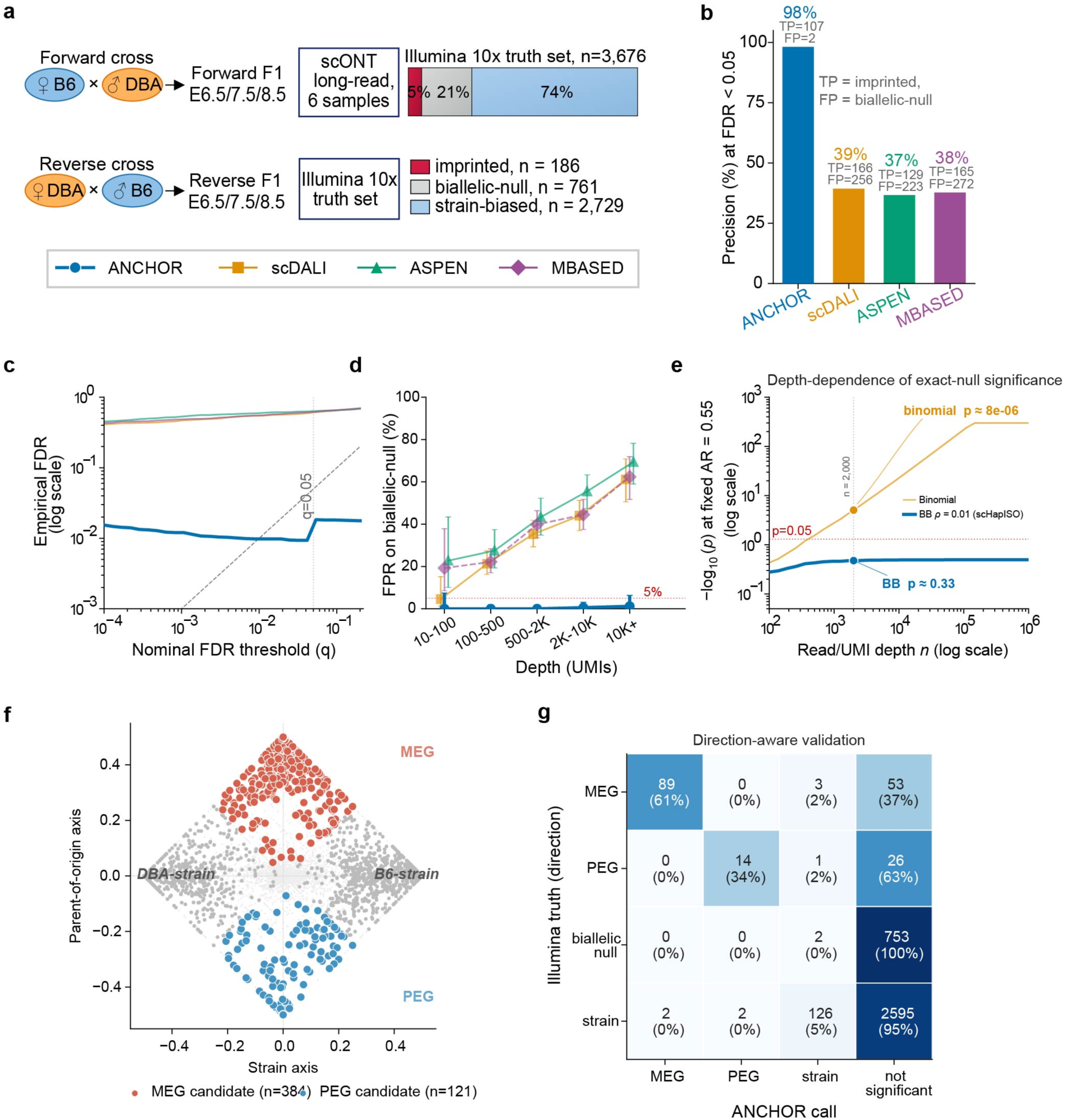
| Allele-specific-expression benchmark on single-cell long-read F1 embryos using an independent biallelic negative-control set. (**a**) Reciprocal hybrid (C57BL/6J × DBA/2 F1) mouse gastrulating embryos (E6.5/7.5/8.5; six scONT libraries); independent Illumina truth set of 3,676 genes (186 imprinted, 761 biallelic-null, 2,729 strain-biased). (**b**) At each method’s native FDR < 0.05, imprinted true-positive and biallelic-null false-positive calls for ANCHOR, scDALI-Hom, ASPEN and MBASED. Imprinted true positives (of 186): 107, 166, 129 and 165; biallelic-null false positives: 2/759, 256/739, 223/473 and 272/717 on their respective method-native testable sets. ANCHOR has the highest specificity and the lowest imprinted recall — a specificity–sensitivity trade-off (see Extended Data Fig. 8b–d for an effect-size-matched comparison). (**c**) Empirical FDR versus nominal Benjamini–Hochberg *q* (log–log); dashed, *y* = *x*. (**d**) False-positive fraction on biallelic-null controls across five UMI-depth bins (Wilson 95% confidence intervals); dotted, 5%. (**e**) Analytical −log₁₀(*P*) for a binomial versus a Beta-binomial test (ρ = 0.01) at allelic ratio 0.52 versus UMI depth. (**f**) Reciprocal-cross scatter rotated 45° so the strain axis (Δ_F + Δ_R)/2 is plotted against the parent-of-origin axis (Δ_F − Δ_R)/2, where Δ = AR − 0.5 and subscripts F and R denote the forward and reverse cross; maternally (red, *n* = 127) and paternally (blue, *n* = 121) expressed candidates are autosomal genome-wide discovery-space calls, not restricted to the 186 reference genes; the X chromosome is excluded throughout the parent-of-origin analysis (Fig. 5). (**g**) Direction-aware confusion of Illumina truth against ANCHOR call (n = 186 imprinted truth genes) over four call categories (maternal, paternal, strain-biased, not-called); count and row fraction (per-direction sensitivity) per cell. Of 186, 122 genes were called (113 correct direction) and 64 not called.

The cause is a depth-dependent calibration failure: with thousands of informative UMIs, an exact-null test turns a biologically negligible allelic-ratio deviation into an overwhelmingly significant P value (Fig. 4c–e), whereas a fixed Beta-binomial overdispersion (ρ = 0.01) keeps such deviations non-significant (Fig. 4e). The benchmark therefore measures detection of robust allelic imbalance rather than any statistically measurable departure from a 0.5 ratio. The high-depth failure mode is what the negative-control set exposes: shared false-positive characterization in Extended Data Fig. 8e–h and per-gene effect-size distributions in Extended Data Fig. 8i–k. The overdispersion ρ = 0.01 was fixed before this benchmark from matched-Illumina technical dispersion and was not tuned on the biallelic-null genes. Across ρ = 0.0025–0.05 the Beta-binomial still keeps these deviations non-significant (Extended Data Fig. 8l,m), so the specificity gain reflects the model rather than a value of ρ chosen on this benchmark.

Projected onto rotated strain and parent-of-origin axes, ANCHOR’s autosomal genome-wide calls resolve into 127 maternally and 121 paternally biased candidates (Fig. 4f). Beyond rejecting false positives, ANCHOR assigns parent-of-origin *direction* correctly: of the 186 imprinted truth genes, it assigned a direction to 122 (a broader set than the 107 reaching FDR < 0.05) and left 64 uncalled. Of these 122, 113 were correct and 1 was inverted, a strict precision of 113/122 (93%) that counts direction-flipped and strain mis-calls as errors (**Fig. 4g**). Together, these results show that ANCHOR reaches near-zero false positives on biallelic-null controls without a tunable effect-size threshold while assigning parent-of-origin direction at 93% precision, trading sensitivity for specificity on robust-effect ASE.

### ANCHOR maps a cell-type-resolved parent-of-origin imprinting landscape in gastrulation

We next used reciprocal-cross imprinting as a stringent biological stress test, because resolving it requires all three ANCHOR layers together: de novo informative-site recovery, molecule-level allele assignment and isoform-resolved quantification. ANCHOR learns only the two strain haplotypes of the supplied diploid sample, so parent-of-origin assignment is a downstream mapping from the reciprocal crosses and never enters the evidence model. Decomposing each gene’s forward and reverse allelic bias into a strain-invariant *cis* component, (*b*∼F∼ + *b*∼R∼)/2, and a parent-of-origin component, (*b*∼F∼ − *b*∼R∼)/2, separates imprinting from strain effects genome-wide (Extended Data Fig. 9a–c), robust across minimum read coverage (Extended Data Fig. 9d,e). As a qualitative check on the decomposition, X-linked genes show strong parent-of-origin signal in extra-embryonic lineages (median |POI| 0.469 versus 0.036 for autosomal genes; gene-by-stage observations, n = 170 versus 21,752; one-sided Mann–Whitney *U*, *P* ≈ 9.4 × 10^−88^, reported descriptively because stage replicates of a gene are not independent; Extended Data Fig. 9f,g), consistent with the expected maternal-X bias of extra-embryonic tissues^25^. This maternal-X signal reflects both imprinted X-inactivation in female embryos and the single maternal X of every male cell in these sex-pooled libraries, and Y-linked expression was insufficient to assign sex per cell. We therefore do not use X-linked genes to discover loci and treat the X-linked pattern only as a qualitative positive-control check on reciprocal-cross decomposition, not as a calibrated control or a sex-composition, mapping or dosage readout. The decomposition’s specificity is instead anchored on recovered canonical autosomal imprinted genes. We make no novel X-linked imprinting claims.

Requiring reciprocal direction inversion, reproducibility across ≥ 2 developmental stages and recovery of known controls, ANCHOR defines a 29-gene cell-type-specific parent-of-origin imprinting (CT-POI) panel: 22 established imprinted genes (16 classical, 5 non-canonical extra-embryonic25 and the isoform-conflict gene H13) and 7 reciprocal-cross-validated candidate genes not catalogued as canonical imprinted, comprising 21 paternally and 8 maternally expressed genes retained under the pre-specified reciprocal-inversion and ≥2-stage reproducibility rules (**Fig. 5a,b**; Supplementary Fig. 1, 2). Because each cross-stage combination is a single pooled library (Methods), the ≥2-stage criterion is cross-stage reproducibility across distinct developmental conditions rather than within-condition biological replication. The complete gene-by-cell-type evidence (forward/reverse allelic ratio, informative UMIs per cross and stage, expressing cells, POI and *cis* effect, FDR *q*, class and reproducibility status) is in Supplementary Table S6. Every retained CT-POI gene shows the pre-specified reciprocal direction-inversion pattern and lies closer to the parent-of-origin than the strain axis of the reciprocal-cross scatter (**Fig. 5c**), and imprinting is cell-type-restricted rather than uniform: *Jade1*, for example, is expressed in 33 cell types but paternally imprinted in only three extra-embryonic types (Fig. 5d; Extended Data Fig. 7b), and *H13* is maternally imprinted in 16 of 36 cell types (Fig. 5d).

**Fig. 5.**
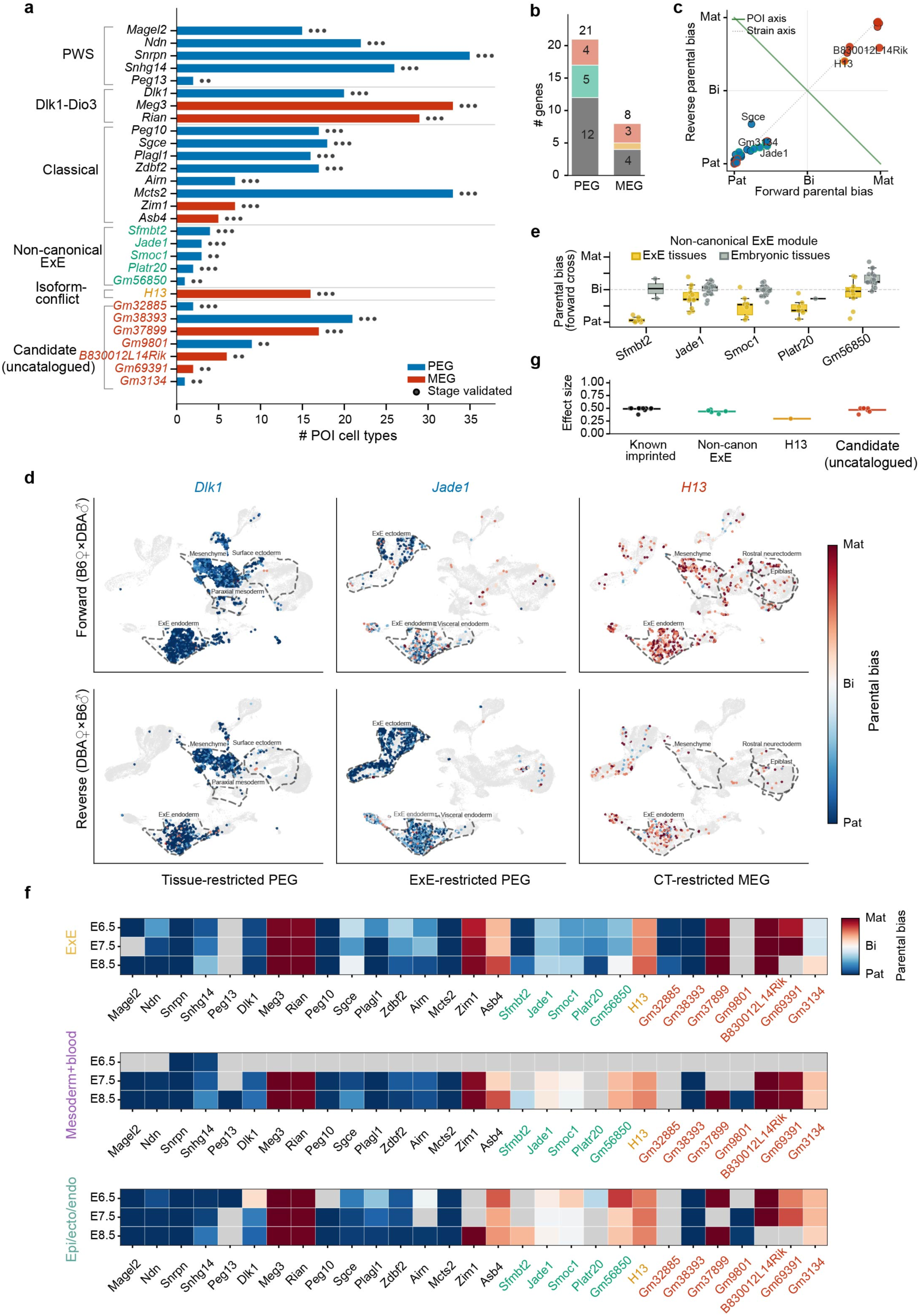
| Cell-type-specific parent-of-origin imprinting (CT-POI) landscape in mouse gastrulation. (**a**) Census of 29 CT-POI genes from hybrid mouse gastrulating embryos profiled by single-cell ONT, including 22 established imprinted genes and 7 reciprocal-cross-validated candidates not catalogued as canonical imprinted genes (several correspond to documented non-canonical imprinted loci, see Supplementary Table S6). Genes are grouped by module; bar length indicates the number of POI-positive cell types. (**b**) Stacked paternal/maternal totals by category. (**c**) Reciprocal-cross scatter; *x*, forward bias; *y*, reverse bias; green line, parent-of-origin axis; grey dotted, strain axis. (**d**) Paired UMAPs (forward top, reverse bottom) of three genes (*Dlk1*, *Jade1*, *H13*) coloured by per-cell parental-bias score (blue paternal to red maternal); UMI < 4 in grey; dashed contours, POI cell types with ≥ 10 expressing cells. (**e**) Per-cell-type parental bias (forward) for five non-canonical extra-embryonic genes, extra-embryonic versus embryonic. (**f**) Lineage × stage × gene heatmap of pseudobulk parental bias (forward); three lineage sub-heatmaps; light grey, < 10 UMIs; white, expressed but biallelic. (**g**) Per-gene maximum effect size by category.

Imprinting concentrates in extra-embryonic endoderm and ectoderm and resolves into two regulatory modes: broadly imprinted clusters (Prader-Willi/Angelman26, Dlk1–Dio327) active across lineages, versus extra-embryonic-restricted loci (Fig. 5e,f; Extended Data Fig. 9h–j). Per-gene maximum effect sizes are shown by imprinting category (Fig. 5g).

At isoform resolution, ANCHOR exposes imprinting that gene-level counting averages away (Extended Data Fig. 10a). UMIs without a recoverable junction chain are retained as an explicit unassigned category rather than discarded (per-sample isoform-assignable and unassigned UMI counts are in Supplementary Table S7). Across the 84,547 isoform-assignable gene, isoform and cell-type combinations (Extended Data Fig. 10b), isoform-resolved analysis recovers parent-of-origin signal that gene-level counting can mask or dilute, with hybrid-validated events spanning multiple cell types (Extended Data Fig. 10c). For example, H13 is biallelic at the gene level yet maternally biased at the isoform level (Extended Data Fig. 9k,l; Extended Data Fig. 10d), and Snhg14 and Snrpn show the same isoform-level imprinting (Extended Data Fig. 10e,f). The H13 result recovers an isoform-level imprinting effect first reported in bulk tissue28, providing a positive control. Together, these results show that ANCHOR resolves a 29-gene cell-type-restricted parent-of-origin imprinting landscape in gastrulation, separating imprinting from strain effects by reciprocal direction inversion and exposing isoform-level imprinting that gene-level counting masks.

### *Sgce* co-expresses a paternal canonical and a maternal transposon-derived isoform within single cells

*Sgce*, a canonical paternally expressed gene^26^, is paternally biased across its gene body in both crosses (parental-bias score ≈ −0.97; Fig. 6a), consistent with external reciprocal-cross RNA-seq^17,18^. However, ANCHOR additionally detects, from the maternal allele, a truncated isoform whose first detected junction originates within an intronic L1MC1 LINE element^27^: reads spanning the transposon-internal 5′ splice site to the canonical acceptor (Fig. 6b) are maternally biased (parental-bias ≈ +0.52, ∼76% maternal reads) in both crosses (Fig. 6c), and the same junction is recovered at proportionate depth in the matched Illumina data (long-read 4,001 vs short-read 88 reads; ratios 0.90 vs 0.86), excluding a long-read artefact (Fig. 6d; Supplementary Fig. 3). A UMAP gallery resolves the parental bias and expression of the canonical and truncated isoforms across cell types (**Fig. 6e**). At single-cell resolution, 223 of 287 cells expressing both isoforms (77.7%) simultaneously read out a paternal canonical and a maternal transposon-derived isoform (Fig. 6f), the allele-autonomous pattern expected when each parental allele expresses its own isoform. These dual-isoform cells are enriched in extraembryonic lineages (allantois, visceral endoderm and extraembryonic endoderm; Fig. 6e), the tissues that form the maternal–fetal interface. Allele-resolved methylation of the L1MC1 element in external blastocyst PBAT data is lower on the maternal allele (43.5% vs 92.3%; one-sided Fisher *P* = 0.0041; **Fig. 6g**), and AlphaFold-3^30,31^ modelling of the sarcoglycan complex shows a reduced SGCE–sarcoglycan inter-chain iPTM for the truncated isoform (canonical 0.56 vs truncated 0.38; Δ ≈ −0.18) while sarcoglycan core contacts are unchanged (**Fig. 6h**). These observations support an L1MC1-associated alternative-promoter or upstream-exon model in which the maternal isoform would yield a non-assembling subunit, but do not establish transcription-start-site usage, promoter activity or causality; read 5′ ends in 10x cDNA do not pinpoint transcription start sites^33^. The kinship-conflict developmental model^34^ (Supplementary Fig. 4) is stated as a hypothesis, not a tested claim. Together, the *Sgce* locus shows that only joint allele-, isoform– and cell-resolution reveals two oppositely imprinted isoforms, a paternal canonical and a maternal transposon-derived isoform co-expressed within single cells, a pattern invisible to gene-level or reference-isoform counting.

**Fig. 6.**
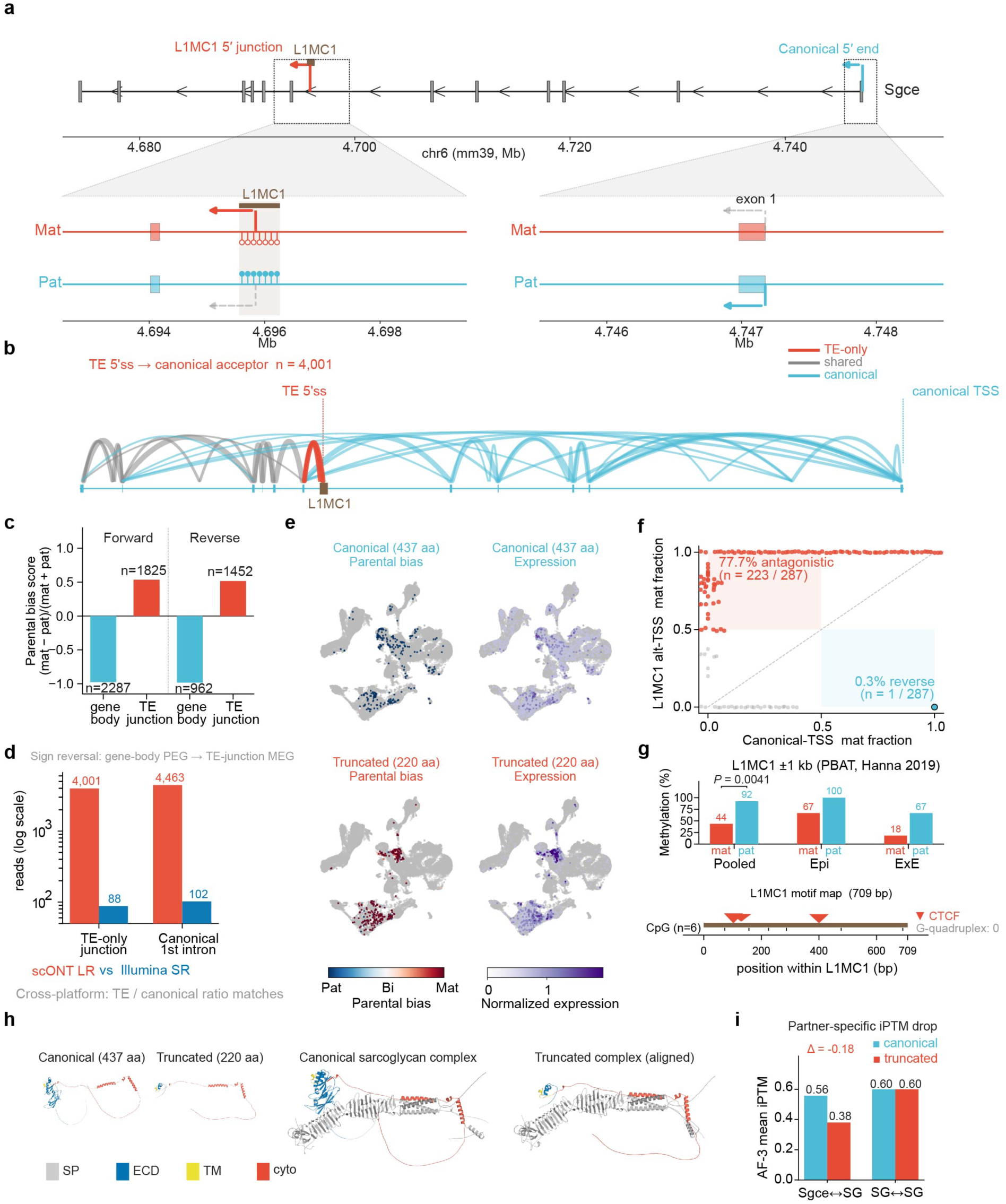
| *Sgce* L1MC1-associated antagonistic imprinting reveals a maternal transposon-junction isoform. (**a**) *Sgce* locus schematic (chr6, mm39): canonical 5′-most exon / first detected junction (blue; not a validated transcription start site) and an L1MC1-associated transposon-junction isoform (red) arising within an L1MC1 element (brown); zoom windows show maternal/paternal tracks with inferred isoform usage and CpG-methylation lollipops. Read 5′ ends in 10x cDNA do not define transcription start sites. (**b**) E8.5 long-read sashimi; arcs coloured transposon-only (red), shared (grey) or canonical-only (blue). (**c**) Parental-bias score for gene-body versus transposon-junction reads in forward and reverse crosses. (**d**) Long-read versus Illumina read counts at the transposon-only junction and the canonical first intron. (**e**) UMAP gallery of parental bias and expression for canonical and truncated *Sgce* isoforms. (**f**) Per-cell scatter (n = 287 cells expressing both isoforms; 223 (77.7%) antagonistic, 1 (0.3%) reverse, the remaining ∼22% concordant or weakly biased) of the L1MC1-associated junction-isoform versus canonical maternal fraction; shaded antagonistic and reverse quadrants. (**g**) Allele-resolved L1MC1 CpG methylation (PBAT) across blastocyst lineages (maternal 43.5% vs paternal 92.3%; one-sided Fisher exact test, *P* = 0.0041), with the L1MC1 motif map (CTCF hits, CpG positions). (**h**) AlphaFold-3 models by SGCE domain: canonical and truncated monomers, four-chain SGCE–sarcoglycan complexes, and inter-chain iPTM (single AlphaFold-3 model_0 per complex, n = 1 per condition, no statistical test) for SGCE–sarcoglycan (canonical 0.557 vs truncated 0.380; with DAG1 0.547 vs 0.380) versus a sarcoglycan–sarcoglycan control (Δ ≈ 0). The truncated isoform lacks the interface-forming extracellular domain, so the lower iPTM is expected and is shown as supporting context, not proof of mechanism.

## Discussion

We present ANCHOR, a method that enables haplotype-aware, isoform-resolved analysis of allele-specific expression from single-cell long reads without requiring a pre-existing phased or sample-specific genotype. This inference remains contingent on sufficient expressed heterozygosity, mappability and long-read coverage, and on a study design in which allele labels can be assigned. The framework combines RNA-only de novo variant calling with molecule-level probabilistic assignment, resolving each transcript molecule to one of two haplotypes.

Two features are central to its performance. The first is RNA-only de novo expressed-SNP calling. Its front end, a signed-graph two-colouring of the molecule-by-site matrix, is closely related to the minimum-error-correction objective of read-backed haplotype phasing^35,36^, but is applied to RNA and feeds a calibrated gradient-boosted cascade. The calibrated cascade accounted for most of the variant-calling performance, whereas graph residuals improved recall at the high-precision operating point. Its performance was more limited in bulk, high-coverage datasets, where features derived from single-cell structure provided less information.

A second important feature is calibration. By aggregating read evidence within UMIs using a Beta-binomial model with fixed overdispersion, ANCHOR reduces the tendency of high-depth count data to make small technical skews appear significant. This conservatism improves specificity where count-based tests are vulnerable to depth-driven false positives, although it can reduce sensitivity for weak allelic effects.

Several limitations remain. First, under the null, ANCHOR produces slightly conservative rather than perfectly uniform P values. Second, chromosome-Y-free libraries preclude per-cell sex assignment and X-linked imprinting claims. Finally, 10x cDNA read 5′ ends should not be interpreted as transcription start sites.

Biologically, ANCHOR uses reciprocal-cross direction inversion to distinguish parent-of-origin effects from strain-biased cis regulation. This extends a classical genetic design to single-cell and isoform-level resolution, allowing imprinting to be assessed in contexts where single-cross, single-individual or non-reciprocal adult datasets are insufficient. In gastrulating mouse embryos, the resulting parent-of-origin landscape was strongly cell-type-restricted and enriched in extra-embryonic lineages. Isoform-level analysis further revealed imprinting patterns that were masked by gene-level aggregation.

*Sgce* illustrates this added resolution. In addition to the expected paternally biased canonical *Sgce* isoform, ANCHOR identified a maternally biased junction isoform arising within an L1MC1 element. The two isoforms therefore showed opposite parental biases, a pattern that would be difficult to resolve with bulk or allele-agnostic analyses. Because the maternal isoform is predicted to lack the assembly domain and is enriched in extraembryonic lineages, it could act as a parent-of-origin-restricted antagonist of the canonical subunit. In this view, the paternal allele would supply the full-length, assembly-competent protein, whereas the maternal allele would contribute a truncated form that cannot assemble, lowering the dose of functional complex. Such an arrangement matches the kinship-conflict model of genomic imprinting, in which paternally expressed genes tend to promote the transfer of maternal resources to the embryo and maternally expressed genes tend to restrain it. More broadly, because ANCHOR does not require a sample-specific phased genotype and uses parental haplotype references only for allele labelling in the reciprocal-cross setting, it should be applicable to other diploid single-cell long-read datasets with sufficient expressed heterozygosity and coverage. This generalization remains to be tested beyond the mouse scONT and human single-cell PacBio datasets analysed here. By resolving allelic and parent-of-origin effects at isoform resolution from single-cell long reads alone, ANCHOR provides a route to studying allele-specific regulation in systems where sample-specific genotypes are unavailable.

## Methods

### Animals and reciprocal crosses

C57BL/6J (B6) and DBA/2 (DBA) mice were purchased from Beijing Vital River Laboratory Animal Technology Co., Ltd. Reciprocal F1 embryos were generated by crossing the two strains in both directions: forward crosses (B6 ♀ × DBA ♂) and reciprocal crosses (DBA ♀ × B6 ♂). Embryos were collected at embryonic days E6.5, E7.5 and E8.5 from each cross direction, yielding six libraries in total (one per cross × stage). All experimental protocols involving mice were approved by the Ethic Committee of Reproductive Medicine of Reproductive Hospital of Shandong University (2026-58).

### Embryo isolation

Pregnant females were sacrificed and the uterus was transferred to a culture dish containing M2 medium, where adherent blood was thoroughly washed away. Individual decidua-enclosed embryos were dissected out in a clean M2 dish, transferred to a four-well M2 dish and kept on ice. Embryos were washed extensively in 1× PBS to remove maternal blood and red blood spots at the embryo–uterus junction.

### Single-cell dissociation

Embryos were pooled per library according to stage to obtain comparable cell yields: 25-30 embryos for E6.5, 15-20 embryos for E7.5 and 6-8 embryos for E8.5. Pooled embryos were transferred into 200 µl TrypLE Express (Gibco) and mechanically disrupted with forceps into at least two fragments, then triturated ten times with a 200-µl pipette and low-retention tips. The digest was transferred to a 1.5-ml low-binding tube; the dish was rinsed with a further 200 µl TrypLE Express that was added to the same tube, mixed by ten gentle pipetting strokes, and incubated for 3 min at 37 °C in a shaking incubator at 1,000 rpm. The suspension was triturated ten times, incubated for a further 2 min at 37 °C with shaking, and triturated ten times again. Digestion was quenched with 1 ml ice-cold 10% (v/v) FBS in PBS and mixed by five gentle pipetting strokes. Cells were filtered through a 40-µm Flowmi cell strainer. The filtrate was centrifuged at 300g for 5 min at 4 °C and the supernatant was aspirated from the opposite side of the pellet, leaving ∼10 µl to avoid cell loss. Cells were washed once with 1 ml PBS containing 2% FBS by gentle inversion (no pipetting) and centrifuged again to remove the supernatant. The pellet was resuspended in 60 µl PBS containing 2% FBS by approximately five gentle pipetting strokes with low-retention tips. Cell concentration and viability were assessed and adjusted for loading.

### Chromium 10x capture and sibling multi-omics split

Single-cell suspensions were loaded onto the 10x Genomics Chromium platform^37^ targeting ∼6,000 cells per library, following the manufacturer’s instructions for the Chromium Single Cell 3′ Kit (v3). Barcoded full-length cDNA was amplified and then split into two portions for matched short– and long-read sequencing of the same cells (sibling multi-omics), so that the Illumina and Nanopore libraries share cell barcodes (Illumina barcodes carry a –1 suffix).

### Short-read (Illumina) library and sequencing

One portion of the amplified cDNA was processed into a short-read sequencing library following the standard 10x Genomics protocol and sequenced on an Illumina NovaSeq X system (paired-end, 150 bp) at a minimum depth of 20,000 reads per cell, yielding FASTQ output.

### Long-read (Oxford Nanopore) library and sequencing

The matched portion of the amplified cDNA was processed into a long-read library using the Oxford Nanopore Ligation Sequencing Kit V14 (SQK-LSK114) and sequenced on a PromethION 48 system (R10.4.1 chemistry, FLO-PRO114M flow cell; Wuhan Benagen Technology Co., Ltd, Wuhan, China) at a minimum depth of 20,000 reads per cell.

### Basecalling

Raw POD5 files from each Nanopore run were basecalled with Dorado (v1.1) using the super-accurate (SUP) model (v5.0.0), yielding FASTQ output.

### Reference and marker panel

Reads were aligned^38^ to GRCm39/mm39 (10x reference refdata-gex-GRCm39-2024-A)^39^; alignments were sorted and indexed with SAMtools^40^. The DBA/2-versus-B6 marker panel was derived from the Mouse Genomes Project (REL-2112-v8)^41,42^ as homozygous differences at QUAL ≥ 30: 5,569,606 SNPs and 1,182,310 indels (6,751,916 total). scONT reads were processed with wf-single-cell to obtain cell barcodes (CB), UMIs (UB), gene (GN) and transcript (TR) tags; TR tags were not used as isoform ground truth because reference mapping biases allele-specific spliced reads (Supplementary Note 1).

### De novo variant calling (ANCHOR caller)

For each chromosome, per-UMI consensus reads were piled into a molecule-by-site sign matrix *A* ∈ {+1, −1, NaN} (reference/alternate/uncovered). Each molecule and each site was assigned one of two signs (a rank-1 *two-colouring* of the molecule-by-site graph), approximating A ≈ hδ^T^ + R (where h is a per-molecule haplotype sign, δ a per-site phase sign and R the residual), by spectral initialization followed by alternating sign optimization (eight iterations). Each site also received a *frustration* score (the weighted fraction of its observations that disagree with this two-sign assignment) and a graph-negative-mass score (the same disagreement, un-normalized). Thirty-four features fed a two-stage cost-sensitive LightGBM cascade^43^: stage 1 used BAM and context features and stage 2 was a graph-aware reranker, together emitting a phased VCF^44^ with GQ and phase-set fields. The features comprised read-level BAM statistics, derived quality and concordance, single-cell support counts, graph/linkage features, sequence context (including RNA-editing/ADAR^45^ context from REDIportal^46^), and haplotype Bayes factors. Models were trained on chr1–10 (human HG002) or per-leave-one-chromosome-out (mouse) and evaluated on held-out chromosomes. Feature attribution used TreeSHAP^47^. The complete algorithmic specification is provided in Supplementary Note 1, with the full feature list and importances in Supplementary Table S4. It covers the factorization objective and entry weighting; treatment of missing (NaN) entries; spectral initialization; the *h* and *δ* update equations; the convergence criterion; handling of disconnected graph components and phase-set boundaries; indel representation; computational complexity and memory strategy; the exact 34 features; LightGBM hyperparameters; the train/validation/test split and leakage controls; whether calibration is per-chromosome, per-sample or global; and the operating thresholds on the stage-2 classifier probability s2 (production s2 ≥ 0.70; high-precision s2 ≥ 0.95).

### Variant-calling benchmark

Truth sets were GIAB v4.2.1 for human samples. To avoid penalizing callers for sites with no RNA coverage, “callable truth" was defined per coverage threshold as GIAB SNPs intersected with BAM coverage ≥ N (MAPQ ≥ 5, BQ ≥ 10), independently of any caller. Competitors (Clair3-RNA^48,49^, LongCallR^14^) were run with their recommended RNA presets on the identical BAMs and evaluated PASS-only. For het-only/ASE comparisons, competitor calls were restricted to heterozygous SNPs within confident regions to avoid scope-mismatch artefacts. For the same-sample HG002 evaluation (Fig. 2c–e), all evaluated methods were scored on a single shared candidate universe. Generating candidate sites by pileup thresholds is the standard first step of long-read RNA variant callers (for example, LongCallR14 selects candidates at depth ≥ 6, alternate-allele fraction ≥ 0.1 and alternate count ≥ 2). ANCHOR’s candidate universe, the union of per-UMI pileup sites on the held-out chr11–22 (861,717 sites with ≥ 2 alternate UMIs), is comparably liberal. GIAB truth and every caller’s PASS output were clipped to it so that precision and recall share one denominator. The coverage-based callable-truth definition above is used for the cross-sample bulk Iso-Seq comparisons (Fig. 2f–i). chr6 (HLA/MHC) was excluded identically for all methods and truth sets (pair-HMM memory cost in the allelic engine; unreliable truth in the hypervariable MHC), so the exclusion does not advantage ANCHOR.

### Allelic deconvolution (ANCHOR allelic engine)

Per informative site, a pair-HMM^50^ scored each read against the two haplotype reference sequences by global forward dynamic programming, yielding a per-site LLR; LLRs were calibrated by linear (Platt^51^) logistic regression with batch-aware covariates. A read-level HMM integrated calibrated LLRs along the molecule (allele-1 / allele-2 / switch states). UMIs were aggregated by a Beta-binomial model into a four-state posterior: allele 1 (B6), allele 2 (DBA), MIX and UNK. A UMI abstained when its posterior fell into the MIX or UNK state. The global overdispersion estimate was ρ ≈ 0.012; production used ρ = 0.01. UMIs were summed per cell, per gene and per isoform. The pair-HMM state space (match, read-insertion, reference-insertion), transition and emission probabilities, base-quality transform, homopolymer and splice-gap handling, per-site LLR definition, and the Platt calibration scope, together with the read-level HMM states, switch penalty, posterior threshold and MIX/UNK rule, are specified in Supplementary Note 1.

### Isoform assignment

Isoforms were assigned from observed splice-junction structure (pair-HMM + HMM inference) rather than reference transcript mapping^52–55^, so allele-specific splicing is resolved rather than collapsed onto the reference isoform (Supplementary Note 1). Isoform-level analyses used the isoform-assignment outputs, not BAM TR tags. UMIs lacking a recoverable junction chain were retained as an explicit unassigned category rather than discarded; per-sample isoform-assignable (transcript-feature) and unassigned-UMI counts are in Supplementary Table S7. The splice-chain clustering rules (junction-coordinate tolerance, minimum supporting reads, transcript-start/end handling, UMI collapsing, isoform equivalence classes, allele-specific junction handling, and the use of reference annotation only to label rather than seed isoforms) are specified in Supplementary Note 1.

### ASE benchmark

An independent truth set was built from the matched Illumina libraries: genes were labelled imprinted (parent-of-origin direction inversion between the forward and reverse cross, reproducible in at least two of three stages), biallelic-null (allelic ratio 0.35–0.65, stage-replicated in both crosses) or strain-biased (same-direction skew in both crosses). ANCHOR (Beta-binomial), scDALI-Hom^22^, ASPEN^23^ and MBASED^24^ were run on the long-read allele counts; false-positive rate was computed on biallelic-null controls and recall on imprinted truth at each method’s native FDR < 0.05. The production overdispersion parameter ρ = 0.01 was fixed before the ASE benchmark from matched-Illumina replicate technical dispersion (the global moment estimate was ρ ≈ 0.012) and was not selected to optimize false-positive rate on the biallelic-null genes; a ρ sweep (0.0025–0.05) is reported as a sensitivity analysis. Competitor versions, exact command lines, references, filters, whether each was given a genotype or phased VCF (none was; all used RNA BAM only), testable-set definitions and the output field used are tabulated in Supplementary Table S5. Fairness audits (common-testable subset, effect-size threshold sweeps, shared false-positive analysis, cell-type deviation, overdispersion ρ) are in Extended Data Fig. 8.

### Cell QC and cell-type annotation

Cells were taken from the wf-single-cell (scONT) and Cell Ranger (Illumina) filtered cell-barcode matrices. Ambient RNA was estimated and corrected with SoupX^56^ on the Illumina matrices (estimated contamination 1.8–3.8% per sample, all < 5%); cells with high mitochondrial fraction (> 20%; Extended Data Fig. 4h), doublets (scDblFinder^57^) and low-quality cells were removed. Cells were integrated with Seurat^58^ and Harmony^59^ (ScaleData on all genes, θ = 2,2,1) and annotated by TransferData from the Pijuan-Sala gastrulation atlas^60,61^ followed by KNN label smoothing (level-2, 36 cell types); annotations failing a neighbour-consistency check were removed. The integrated atlas comprises 103,254 cells (long-read and short-read); of 68,417 scONT cells passing per-cell QC, 55,729 carried a confident cell-type label and were used for all allelic display items. UMAP^62,63^ coordinates and labels were shared across all allelic display items.

### Parent-of-origin analysis

For each gene/isoform × cell type × stage, the B6 read fraction was computed per cross. Parent-of-origin and *cis* effects were separated as POI = (*b*∼F∼ − *b*∼R∼)/2 and cis = (*b*∼F∼ + *b*∼R∼)/2, where *b* = AR − 0.5; effects were called significant when |effect| ≥ 0.10 and a bootstrap^64^/analytical confidence interval excluded 0. For FDR control, two-sided analytical or bootstrap-derived *P* values for the POI effect were submitted to Benjamini–Hochberg correction^65^; the CI exclusion and |effect| ≥ 0.10 threshold were applied as additional effect-size filters. The primary testing unit was gene × cell type × stage; analyses were never pooled across developmental stages. For each stage, Benjamini–Hochberg FDR was controlled separately for the gene-level and isoform-level analyses, in each case over all autosomal gene × cell-type tests passing the ≥ 10 informative-UMI filter (so multiplicity over the 36 cell types is included in the family). The 29-gene CT-POI panel was then built by a stricter, pre-specified inclusion cascade applied per gene: (i) reciprocal direction inversion, (*b*∼F∼)(*b*∼R∼) < 0; (ii) a per-cross effect-size floor |AR − 0.5| > 0.15 in each cross; (iii) Benjamini–Hochberg FDR < 0.05 in both crosses independently; (iv) ≥ 10 informative UMIs per cross; and (v) reproducibility in ≥ 2 developmental stages. Genes meeting the cascade were assigned an evidence tier by the strength of the reciprocal signal. Because this cascade is a conjunction of per-cross thresholds rather than a single combined-effect test, panel inclusion is more conservative than the genome-wide |effect| ≥ 0.10 significance call. Isoform-level tests used a separate gene × isoform × cell type × stage FDR family and were treated as exploratory unless they passed it. All Mann–Whitney *U* tests are two-sided unless stated; the *Sgce* methylation test (Fig. 6g) is one-sided Fisher’s exact as noted. Maternal fraction was derived per cross direction (forward: B6 = maternal; reverse: DBA = maternal) and displayed as the parental-bias score (maternal − paternal)/(maternal + paternal) ∈ [−1, +1]^66^. Manual allele-resolved IGV inspection using ANCHOR read-level posteriors was performed after automated calling as a qualitative audit only and did not alter the pre-specified CT-POI inclusion set. X-linked loci were used solely as an imprinted-X-inactivation positive-control pattern (per-cell sex was not assignable; see main text) and never as a discovery set.

### *Sgce* analysis

Allele-resolved splice junctions, dual-isoform (canonical / transposon-junction) per-cell tables and cross-platform junction counts were computed from the isoform-assignment outputs; L1MC1 allele-specific methylation used blastocyst PBAT data^29^ (GSE114512) and CTCF^67^ motif scanning used JASPAR^68^ MA0139.1; protein consequences used AlphaFold-3^31^ inter-chain iPTM for the sarcoglycan complex with and without DAG1. Read 5′ ends were not interpreted as transcription start sites^33^.

### Statistics and reproducibility

Tests, sample sizes, centre values and error bars are defined in each figure legend. No data were excluded except chr6 (above) and cells failing the QC above. All reported values derive from the source files cited in the figure legends; no values were simulated. The six cross × stage libraries are the biological units; embryos pooled within a library were not genetically demultiplexed to individuals, and the E8.5 libraries derive from 2–3 pooled embryos, so within-condition biological replication is not available. Ambient (maternal decidual or blood) RNA carries the maternal-strain allele and could in principle inflate maternally biased signal and invert with cross direction. We mitigated and quantified this by extensive PBS washing during embryo isolation, doublet/low-quality removal, and SoupX ambient-RNA correction of the Illumina matrices, where estimated contamination was low (1.8–3.8% per sample, all < 5%). Because measured ambient contribution is small and purely paternally expressed genes, which are immune to maternal soup, are recovered with the expected bias, residual ambient confounding of maternally biased calls is expected to be minor. The per-UMI allelic counts are computed at the read level rather than from the SoupX-adjusted expression matrix, so read-level ambient correction or genetic demultiplexing remains a target for future work. The semi-synthetic benchmark (270 BAMs) construction, including the source sample, the parameter grid (SNPs-per-UMI × reads-per-UMI × allelic ratio × overdispersion) and per-UMI truth labelling independent of the ANCHOR emission model, is specified in Supplementary Note 1. Analysis-engine versions, seeds and bootstrap resample counts are recorded in the code repository.

### Use of large language models

A large language model assisted with code drafting, figure-assembly scripting and language editing under author supervision; it did not generate, select or interpret data, and does not meet authorship criteria. All scientific content and conclusions are the authors’ own.

## Data availability

Single-cell ONT and matched Illumina sequencing data for the hybrid mouse embryos reported in this paper are available from the National Genomics Data Center, using accession number CRAxxxxx. Human HG002/HG004/HG005 Iso-Seq data and GIAB truth sets are publicly available (https://downloads.pacbcloud.com/public/dataset/ and https://ftp-trace.ncbi.nlm.nih.gov/ReferenceSamples/giab/release/). The DBA/2 marker panel derives from the Mouse Genomes Project REL-2112-v8. External datasets: blastocyst PBAT GSE114512^29^; external reciprocal-cross RNA-seq^17,18^.

## Code availability

The ANCHOR software tool is available at https://github.com/fzc1997/ANCHOR and the code reproducing all figures and benchmarks at https://github.com/fzc1997/ANCHOR-paper. During peer review the source is provided as password-protected archives; decompress with the password xxx. Both repositories will be released openly upon publication.

## Author contributions

Z.-C.F., S.Y. and C.Z. were responsible for the methodology. Z.-C.F. and S.Y. developed the ANCHOR framework and designed the overall computational strategy. Z.-C.F. implemented the ANCHOR algorithm, constructed the analysis pipeline, performed benchmarking analyses and carried out the major computational analyses. C.Z. conducted reciprocal-cross mouse experiments, embryo collection, single-cell sample preparation and sequencing-related experiments. Y.Y., Y.X. and X.Y. assisted with mouse breeding, embryo collection and sample processing. T.T., P.L., W.C., R.H. and J.C. assisted with data processing, benchmarking, visualization and downstream analyses. Y.L and H.W. provided technical support for single-cell long-read sequencing library preparation and sequencing optimization. X.C. and Y.K. contributed to structural modelling and related analyses. Y.L., Z.-J.C., T.H., K.W. and S.Y. provided resources, scientific guidance and supervision. Z.-C.F. and S.Y. interpreted the data and wrote the manuscript with input from all authors. All authors reviewed, edited and approved the paper.

## Acknowledgements

We are grateful to Wei Li, Chao Liu and Chunchun Wu for their valuable guidance and assistance in embryo collection and manuscript preparation. We thank Yuwen Ke for technical guidance on single-cell long-read sequencing library preparation. We also acknowledge Wuhan Benagen Technology Co., Ltd. for their support in optimizing third-generation sequencing technologies and providing technical advice during experimental workflow development.

## Extended Data figure legends

**Extended Data Fig. 1.**
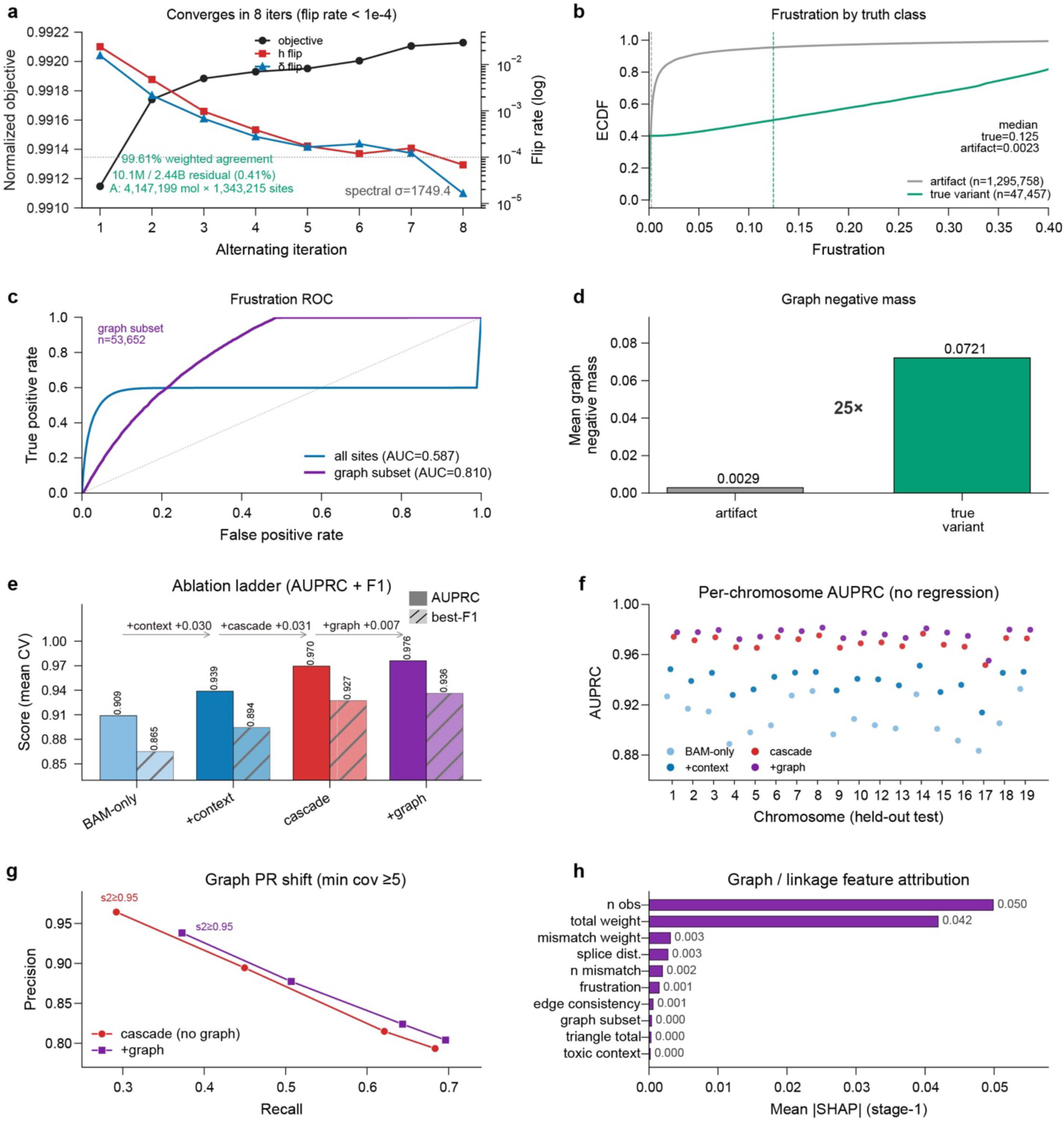
| The ANCHOR caller engine: rank-1 factorization convergence and model-layer contribution. (**a**–**d**) Robust rank-1 diploid factorization on the E7.5 chr11 pilot matrix (4,147,199 molecules × 1,343,215 sites; 2.44 billion entries): (**a**) convergence over eight iterations: the objective and the per-iteration sign-flip rates of the molecule-haplotype vector *h* and site-phase vector *δ* (curves), with the final weighted agreement and residual annotated; (**b**) empirical CDF of the per-site frustration score (the weighted fraction of a site’s molecule signs that disagree with its rank-1 phase assignment) for true variants versus artifacts; (**c**) ROC for frustration over all sites and the linkage-supported subset; (**d**) mean graph negative mass (the same weighted disagreement, un-normalized), artifact versus true variant. (**e**–**h**) Leave-one-chromosome-out cross-validation of four cumulative layers (BAM-only, +context, +cascade, +graph): (**e**) mean AUPRC and best-F1 per layer with incremental gain; (**f**) per-chromosome held-out AUPRC across all 19 autosomes under leave-one-chromosome-out (the single-held-out analysis in Fig. 3e holds out chr11 and reports the remaining 18); (**g**) precision–recall operating points (minimum coverage ≥ 5) for cascade with and without graph features at four thresholds; (**h**) top graph/linkage features by mean |SHAP|.

**Extended Data Fig. 2.**
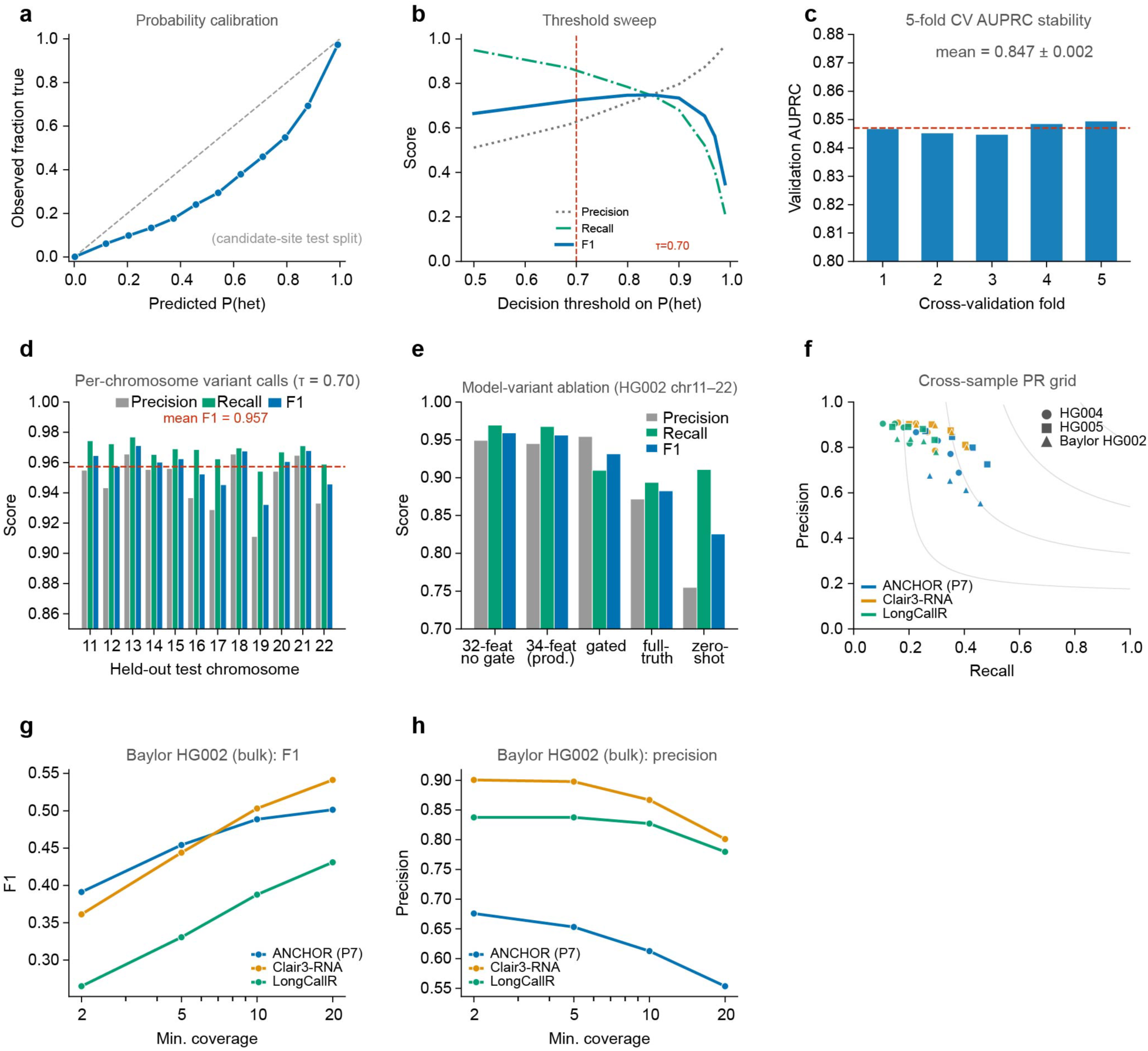
| Reliability, robustness and cross-sample generalization of the ANCHOR (P7) single-cell long-read RNA variant caller. (**a**) Reliability diagram relating the mean predicted P(het) to the observed fraction of true variants across score bins on the HG002 chromosome 11–22 candidate-site test split; the dashed line denotes perfect calibration. (**b**) Precision, recall and F1 of ANCHOR across decision thresholds on P(het) for the same test split; the dashed line marks the production operating point (τ = 0.70). (**c**) Distribution of the validation area under the precision–recall curve (AUPRC) across the five cross-validation folds used in training; the dashed line and shaded band show the mean ± s.d. (**d**) Precision, recall and F1 for each held-out test chromosome (chr11–chr22) at τ = 0.70; the dashed line marks the mean F1. (**e**) Precision, recall and F1 for five ANCHOR model variants on the HG002 test set: the production 34-feature model, a 32-feature model without the null gate, the gated model, a model trained against full (non-callable-restricted) truth, and the mouse-trained model applied zero-shot. (**f**) Precision–recall of ANCHOR, Clair3-RNA and LongCallR across three cross-sample bulk PacBio Iso-Seq datasets (HG004, HG005, Baylor HG002) and four minimum-coverage thresholds (marker shape, sample; colour, method); grey curves are iso-F1 contours. (**g**,**h**) F1 (**g**) and precision (**h**) as a function of the minimum BAM-coverage threshold on the Baylor HG002 bulk dataset for the three callers.

**Extended Data Fig. 3.**
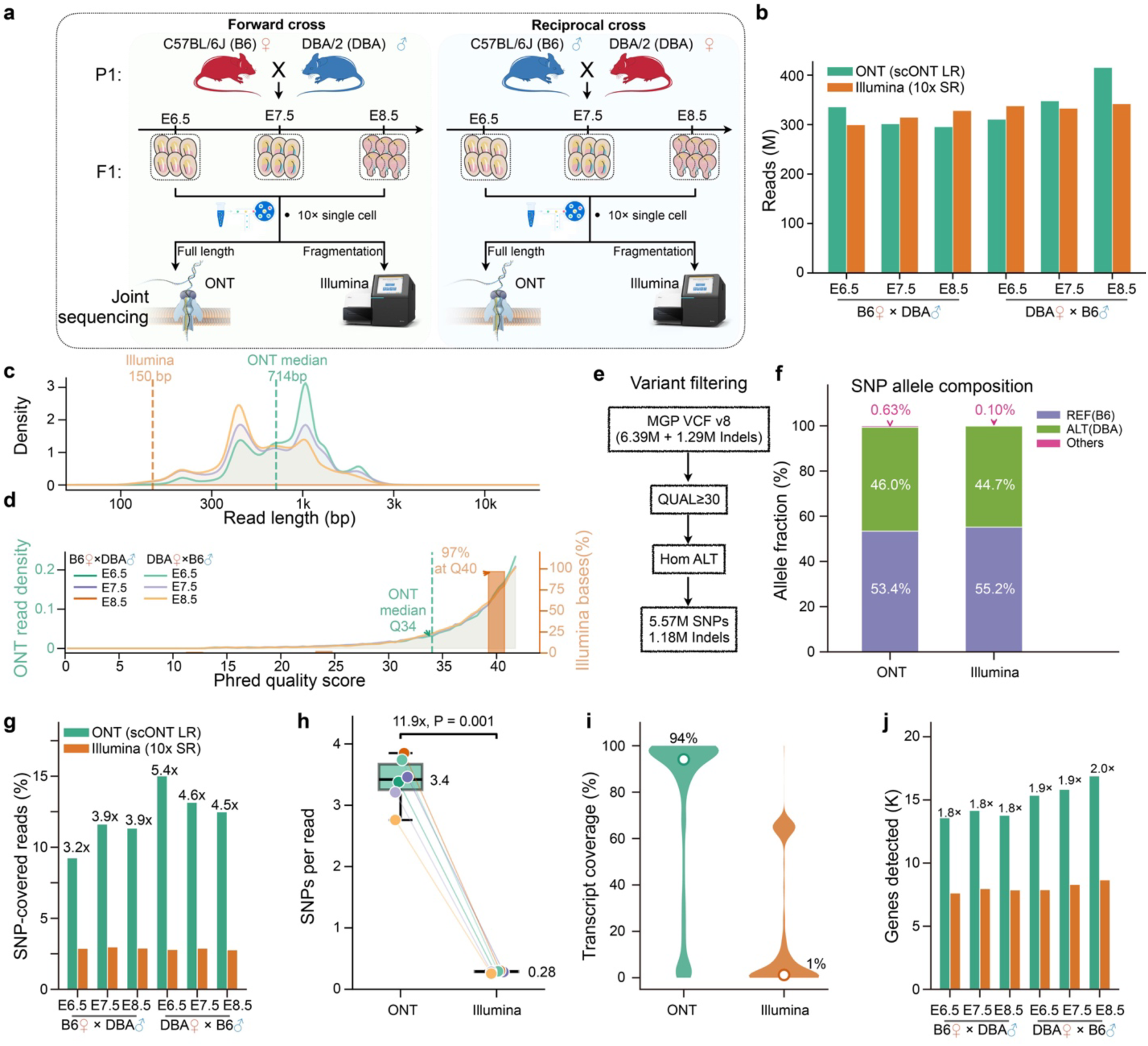
| Data reliability and the long-read advantage of scONT over Illumina. (**a**) Design: reciprocal C57BL/6J × DBA/2 crosses (E6.5/7.5/8.5), profiled by sibling multi-omics (one barcoded cDNA pool split between scONT long reads and Illumina short reads). (**b**) Sequencing yield (percentages, ONT full-length fraction). (**c**) Read-length density (ONT ∼772 bp vs Illumina ∼150 bp). (**d**) Per-read Phred for ONT (median ≈ Q43) and per-base quality at marker SNPs for Illumina (97% at Q40). (**e**) Variant-filtering scheme defining the marker panel (5.57 M SNPs + 1.18 M indels, QUAL ≥ 30). (**f**) Allele composition at marker SNPs (E6.5). (**g**) Percentage of reads covering ≥ 1 marker SNP. (**h**) Marker SNPs per read (Mann–Whitney *U*). (**i**) Per-cell transcript coverage fraction. (**j**) Genes detected per sample. Green, ONT; orange, Illumina.

**Extended Data Fig. 4.**
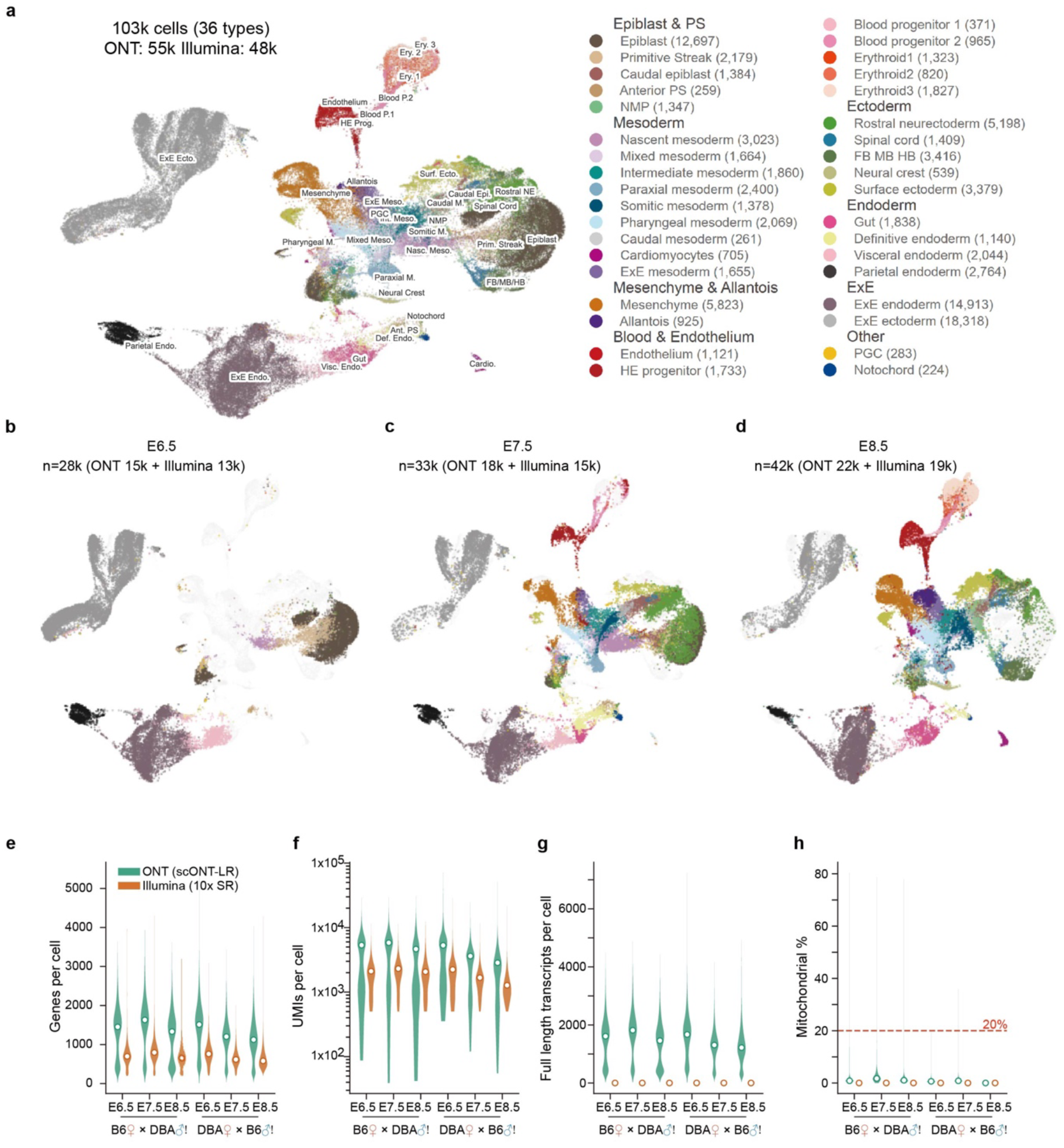
| Joint single-cell atlas and matched per-platform quality control. (**a**) Harmony-integrated UMAP of 103,254 cells (six embryos, both crosses, ONT and Illumina) coloured by cell type (36 types, eight lineages; counts in legend; text marks centroids). (**b**–**d**) The same embedding split by stage (E6.5/7.5/8.5), each combining both crosses; long-read and short-read counts annotated. (**e**–**h**) Per-cell QC as paired violins comparing ONT (teal; n = 68,417 QC-passed cells) and Illumina (orange; n = 57,748); white dot, median: (**e**) genes per cell; (**f**) UMIs per cell (log); (**g**) full-length transcripts per cell (Illumina ≈ 0); (**h**) mitochondrial read percentage (dashed, 20% threshold).

**Extended Data Fig. 5.**
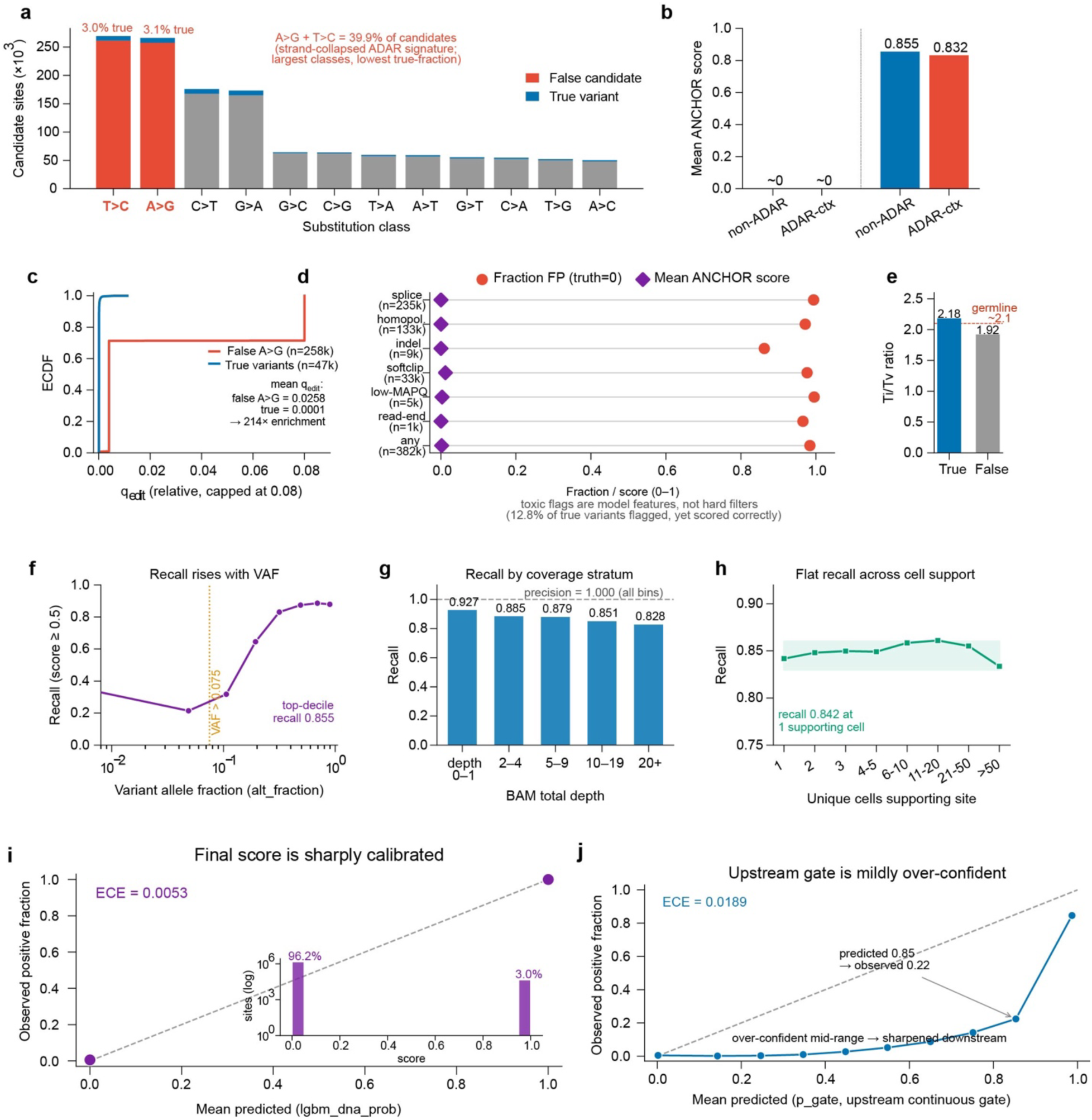
| ANCHOR robustness: RNA-editing and artifact discrimination, stratified recall, and score calibration. All panels derive from the E7.5 chromosome-11 pilot candidate-site set (1,343,215 sites). Panels **a–e** address discrimination of true variants from RNA-editing and technical artifacts. (**a**) Stacked composition of candidate sites by 12-class substitution type, partitioned into true variants and false candidates, with the strand-collapsed ADAR signature (A>G, T>C) highlighted. (**b**) Mean ANCHOR score for false and true sites, split by ADAR sequence context. (**c**) Empirical cumulative distribution of the relative editing score q_edit for false A>G candidates versus true variants. (**d**) For seven artifact-context flags, the fraction of flagged sites that are false alongside the mean ANCHOR score they receive; the flags are model features, not hard filters. (**e**) Transition/transversion ratio for true versus false sites, relative to the germline expectation (2.1). Panels **f–i** characterise performance and calibration. (**f**) Recall as a function of variant allele fraction. (**g**) Recall across BAM coverage strata, with precision noted. (**h**) Recall as a function of the number of unique supporting cells. (**i-j**) Reliability diagrams for the final ANCHOR score (i) and the upstream continuous gate (j), each annotated with its expected calibration error (ECE); the inset in (i) shows the score-distribution histogram. n values are given in each panel.

**Extended Data Fig. 6.**
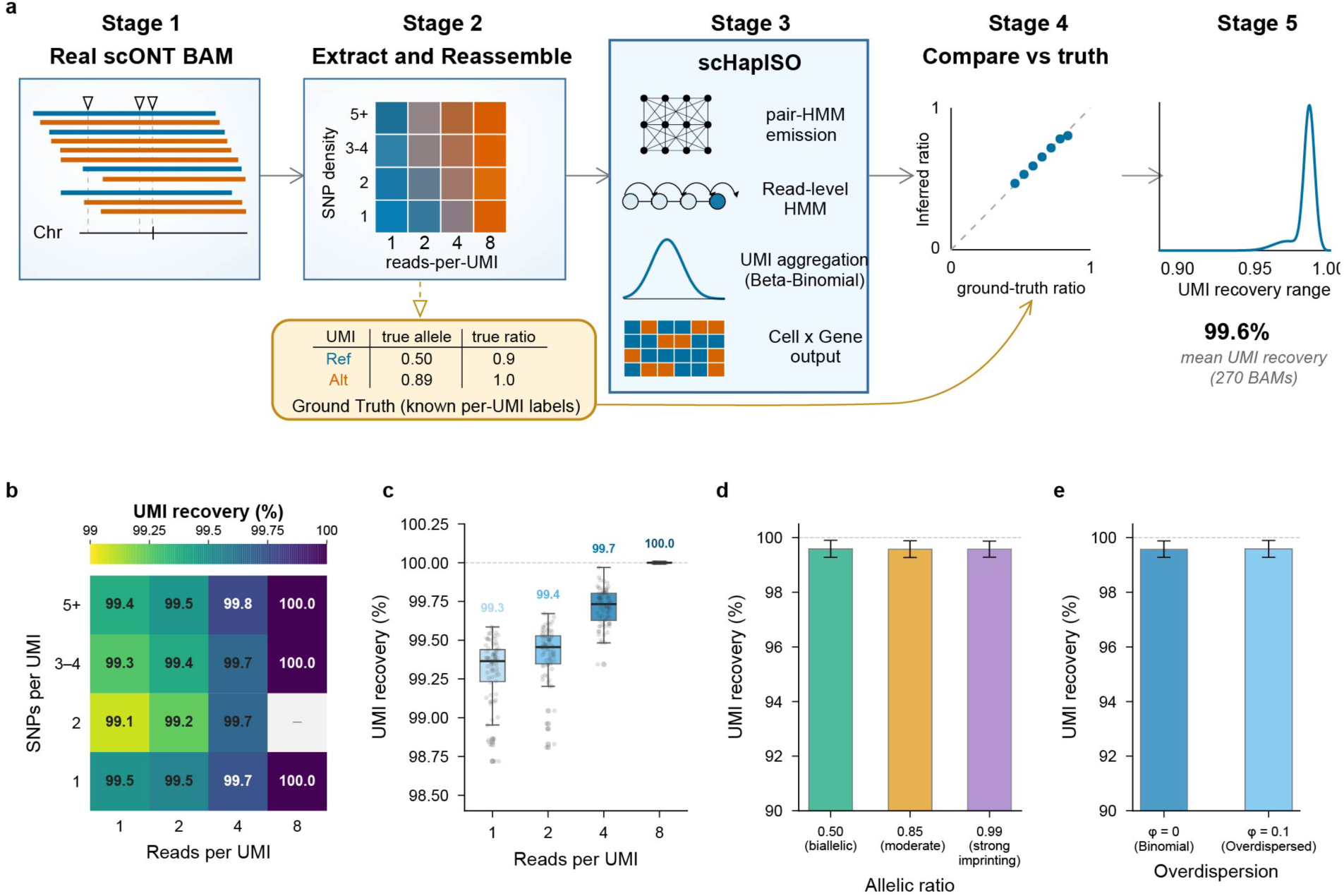
| Semi-synthetic benchmarking of ANCHOR allelic-label recovery across read, variant and dispersion regimes. (**a**) Schematic: reads from real scONT BAMs are reassembled into molecules with known per-UMI allelic labels, processed by ANCHOR, and inferred versus ground-truth allelic ratios compared over 270 synthetic BAMs. (**b**) Mean UMI recovery versus informative SNPs per UMI (rows) and reads per UMI (columns); grey, no data. (**c**) UMI-recovery distribution by reads per UMI (box, median and IQR; whiskers, 1.5× IQR; points, BAMs). (**d**) Mean UMI recovery for allelic ratios 0.50, 0.85 and 0.99 (error bars, s.d.). (**e**) Mean UMI recovery without (φ = 0) and with (φ = 0.1) overdispersion (error bars, s.d.). UMI recovery is the fraction of UMIs whose allelic label matches the synthetic truth (overall 99.58% ± 0.30%).

**Extended Data Fig. 7.**
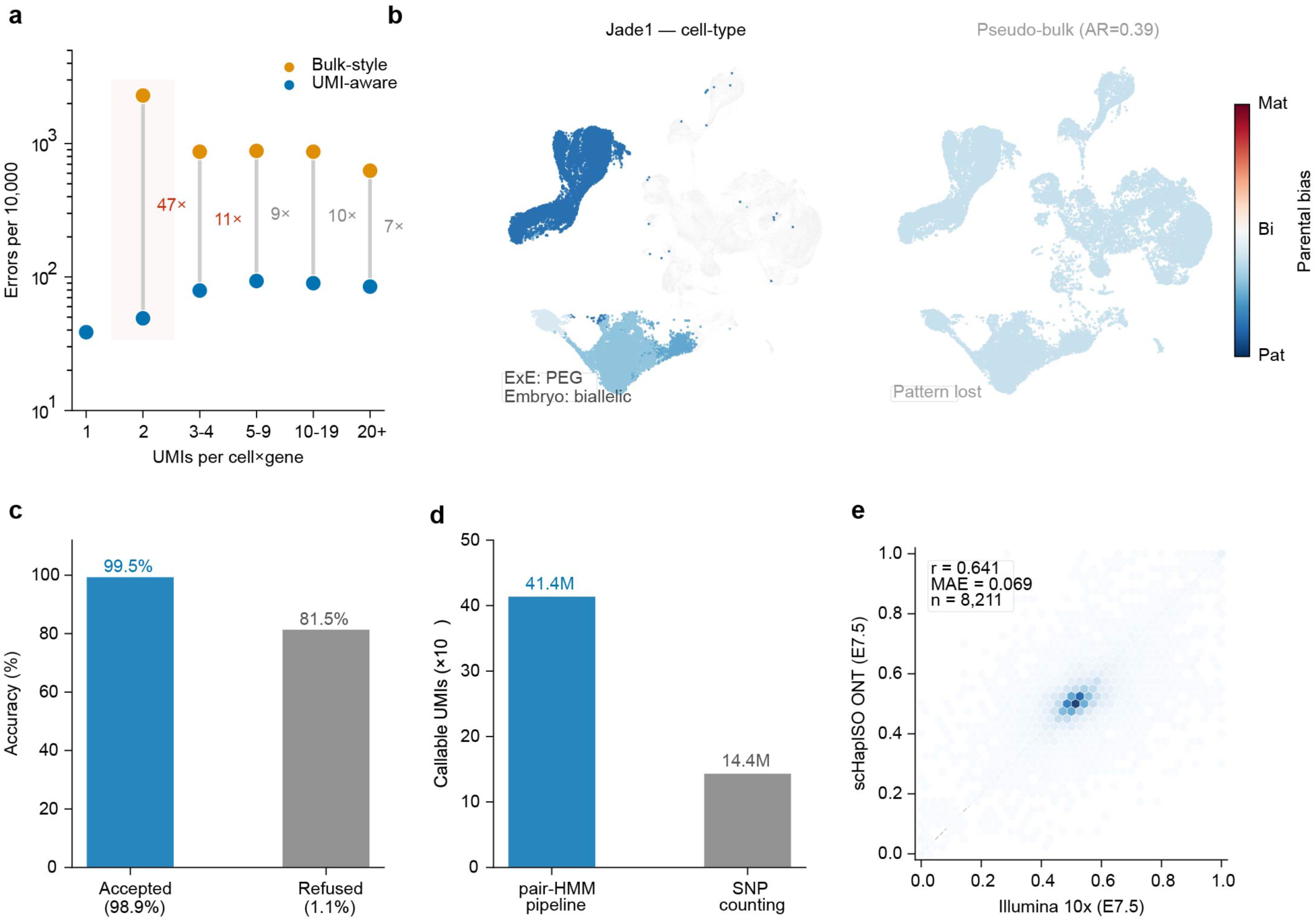
| Per-UMI parental ancestry from single-cell ONT data with ANCHOR. (**a**) Allelic-calling error rates stratified by UMIs per cell × gene, comparing ANCHOR’s UMI-aware Beta-Binomial aggregation (blue) with a bulk-style baseline that pools reads across UMIs (orange). Means over six F1 embryos (forward and reciprocal C57BL/6J × DBA/2 crosses) benchmarked against sibling Illumina 10x calls; grey dumbbells link paired bins, and fold-change labels mark bins with ≥1.5× excess baseline error. (**b**) UMAP of 103,254 merged cells coloured by parental bias for the paternally expressed gene Jade1. Left, ANCHOR per-cell-type bias, with high-coverage extra-embryonic lineages (ExE ectoderm and endoderm, visceral and parietal endoderm) highlighted and embryonic cells with weak bias shown as grey background. Right, the same cells coloured by a single pseudo-bulk allelic ratio. Colour bar, parental bias score (mat − pat)/(mat + pat). (**c**) Accuracy of decidable (B6 or DBA) UMI calls versus the accuracy that abstaining (MIX or UNK) UMIs would have if forced to call; bracketed percentages give each group’s share of all UMIs, averaged across six samples. (**d**) UMIs assignable to a parental haplotype in E6.5, comparing ANCHOR’s pair-HMM emission model (blue) with a pileup-based SNP-counting baseline (grey; HP-tag majority vote over 5.6 M genome-wide homozygous DBA/2 SNPs). (**e**) Hexbin density of pseudo-bulk gene-level B6 allelic ratio in E7.5 between ANCHOR ONT (y) and Illumina 10x (x), for genes with ≥10 allelic UMIs on each platform. Dashed line, y = x; Pearson r, mean absolute error and gene count are inset.

**Extended Data Fig. 8.**
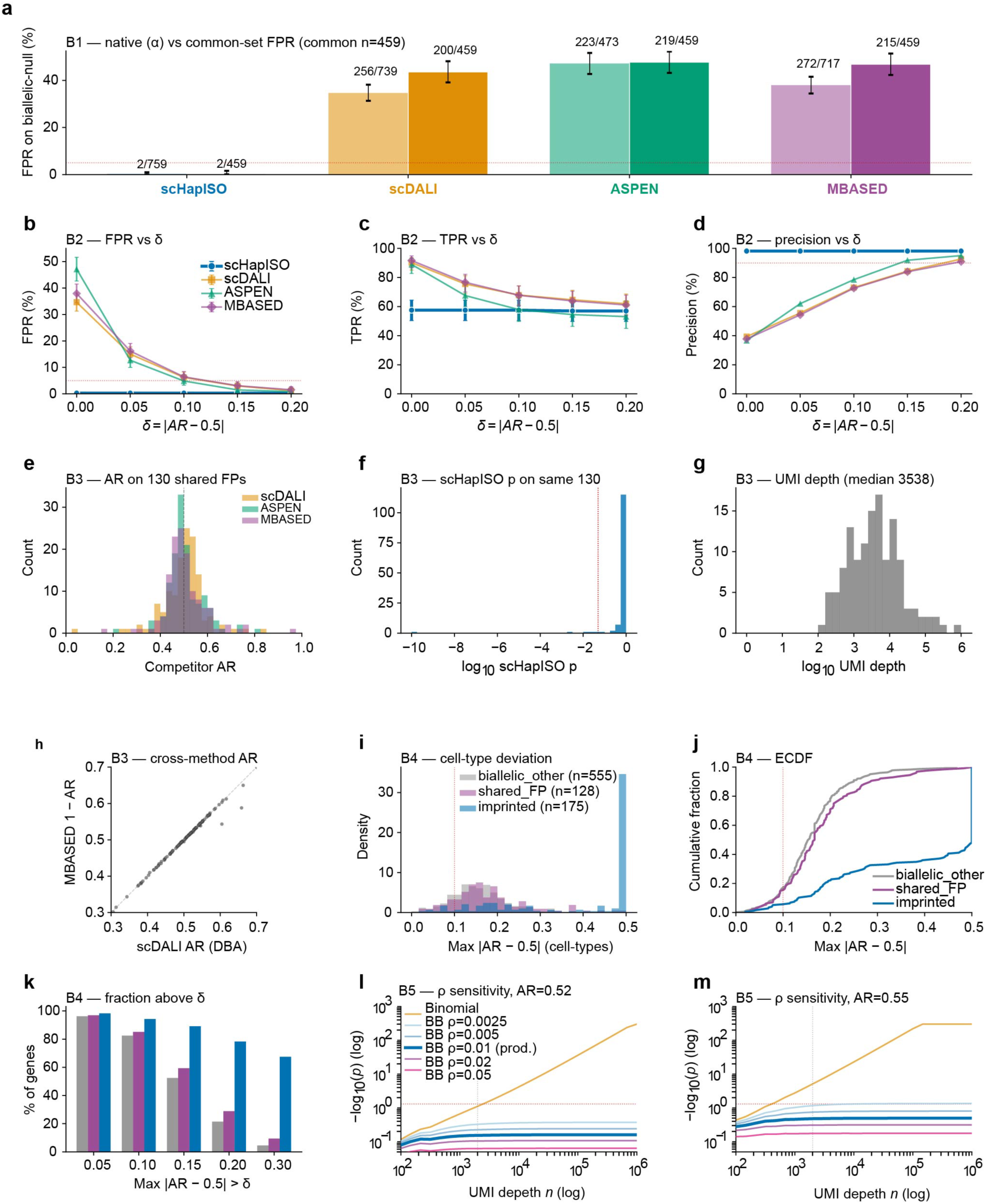
| ASE benchmark fairness audits. (**a**) Per-method false-positive rate on biallelic-null genes at native FDR < 0.05 in each method’s native testable set and in the 459-gene common intersection (Wilson 95% CI; dotted, 5%). (**b**–**d**) False-positive rate (**b**), true-positive rate on imprinted truth (**c**) and precision (**d**) versus effect-size threshold δ = |AR − 0.5| (0–0.20). (**e**) Competitor AR histogram on 130 shared biallelic-null false positives. (**f**) ANCHOR *P*-value and (**g**) UMI depth histograms on the same genes. (**h**) Cross-method AR agreement on the shared false positives. (**i**) Per-gene max |AR − 0.5| density across pseudobulk bins by truth category; (**j**) its empirical CDF; (**k**) fraction exceeding δ thresholds. (**l**,**m**) Analytical −log₁₀(*P*) for binomial versus Beta-binomial tests at ρ ∈ {0.0025–0.05} versus depth at AR = 0.52 (**l**) and 0.55 (**m**); production ρ = 0.01 in bold.

**Extended Data Fig. 9.**
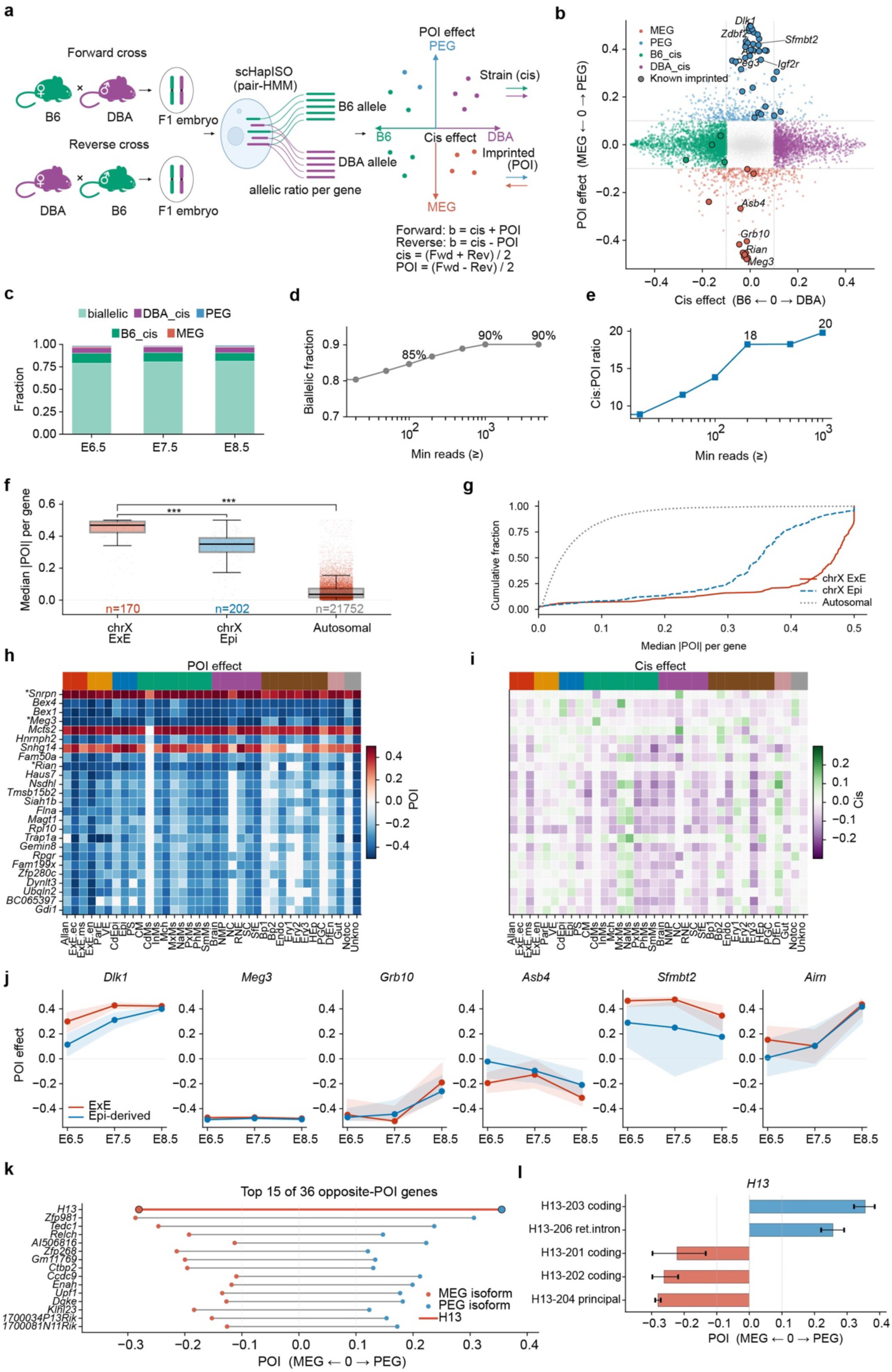
| Reciprocal-cross decomposition of single-cell allelic expression into parent-of-origin and cis-genetic effects during mouse gastrulation. (**a**) Analytical strategy: F1 embryos from forward (C57BL/6J ♀ × DBA/2 ♂) and reverse (DBA/2 ♀ × C57BL/6J ♂) crosses are assayed by single-cell long-read sequencing; per-gene allelic ratios from the two crosses are combined to separate the strain-invariant cis effect from the parent-of-origin (POI) effect. (**b**) Gene-level landscape of cis effect (x; B6 ↔ DBA) against POI effect (y; maternally [MEG] ↔ paternally [PEG] biased); each dot is one gene at one stage, coloured by class, black-outlined dots are known imprinted genes, dashed lines mark the effect-size threshold. (**c**) Class composition per developmental stage. (**d**) Biallelic fraction and (**e**) cis-to-POI ratio as functions of minimum read coverage. (**f**) Per-gene median |POI| for X-linked genes in extra-embryonic (ExE) and epiblast-derived lineages versus autosomal genes (box plots, Mann–Whitney U) and (**g**) the corresponding cumulative distributions. (**h**) Cell-type-resolved POI and (i) cis effects for top-ranked genes, columns ordered by super-lineage (colour bar); asterisks mark known imprinted genes. (**j**) POI effect across stages for six pre-specified imprinted loci in ExE versus epiblast-derived lineages (shading, 95% confidence interval). (**k**) Isoform-level POI for genes carrying oppositely imprinted isoforms (dumbbells span the per-gene isoform POI range) and (**l**) per-isoform POI for H13 (error bars, 95% confidence interval). MEG, rose-red; PEG, steel-blue; B6, green; DBA, purple.

**Extended Data Fig. 10.**
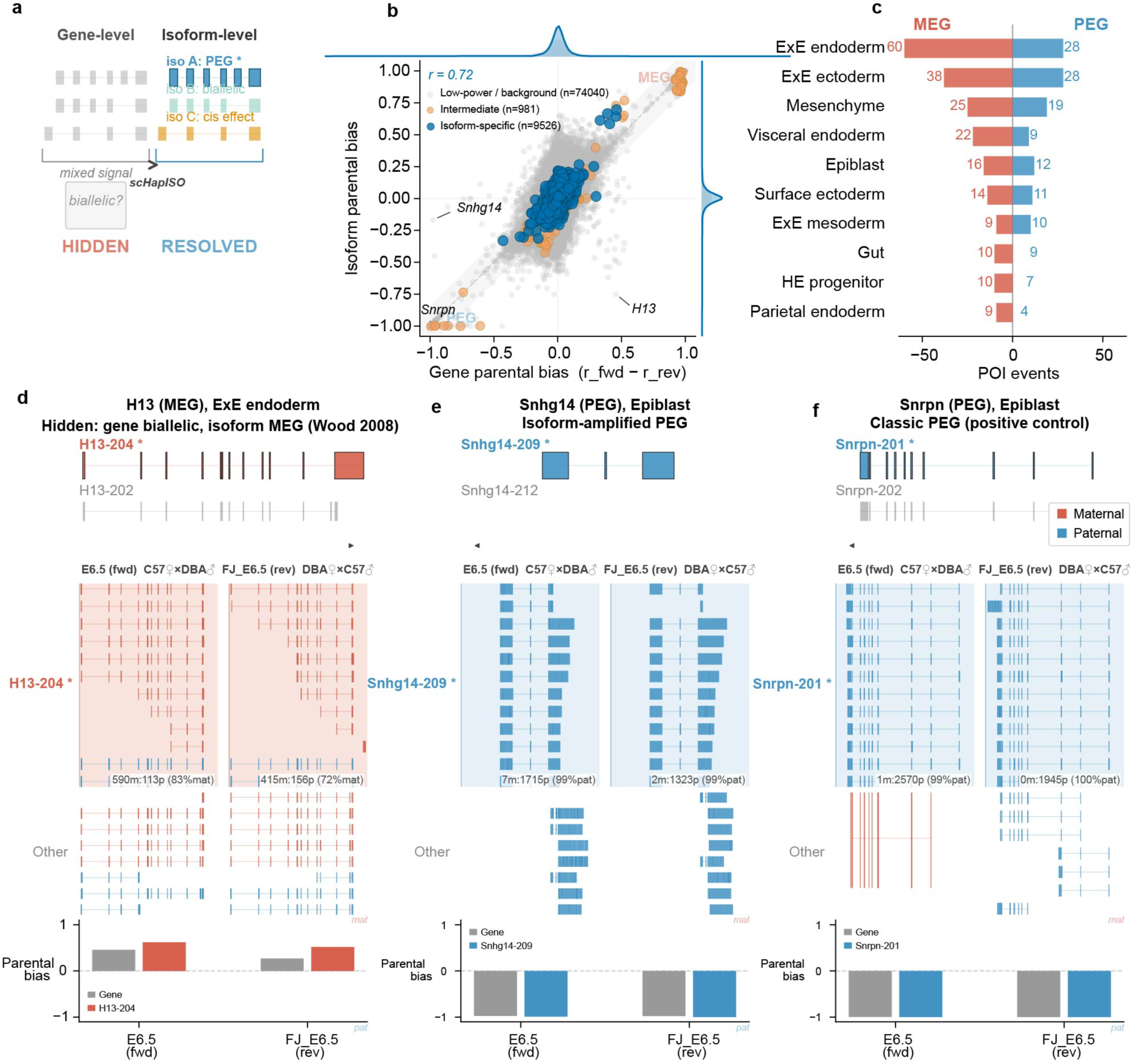
| Isoform-resolution parent-of-origin imprinting in gastrulation. (**a**) Schematic: gene-level quantification averages co-transcribed isoforms (a heterogeneous locus looks biallelic), whereas haplotype-aware isoform-level assignment instead resolves the constituent isoforms separately — here one paternally expressed isoform, one biallelic isoform and one cis-strain-affected isoform. (**b**) Gene versus isoform parental bias for 84,547 gene × isoform × cell-type combinations; grey, low-power; blue, isoform-specific candidates; *r*, Pearson correlation across the isoform-specific subset; three case genes labelled. (**c**) Hybrid-validated parent-of-origin isoform events per cell type (top 10; maternal left, paternal right). (**d**–**f**) Example loci with transcript annotation, allele-tagged read pile-ups in both crosses, and gene-versus-isoform bias: (**d**) *H13* (maternal, extra-embryonic endoderm); (**e**) *Snhg14* (paternal, epiblast); (**f**) *Snrpn* (paternal, epiblast).

## Supplementary information legends

**Supplementary Fig. 1.**
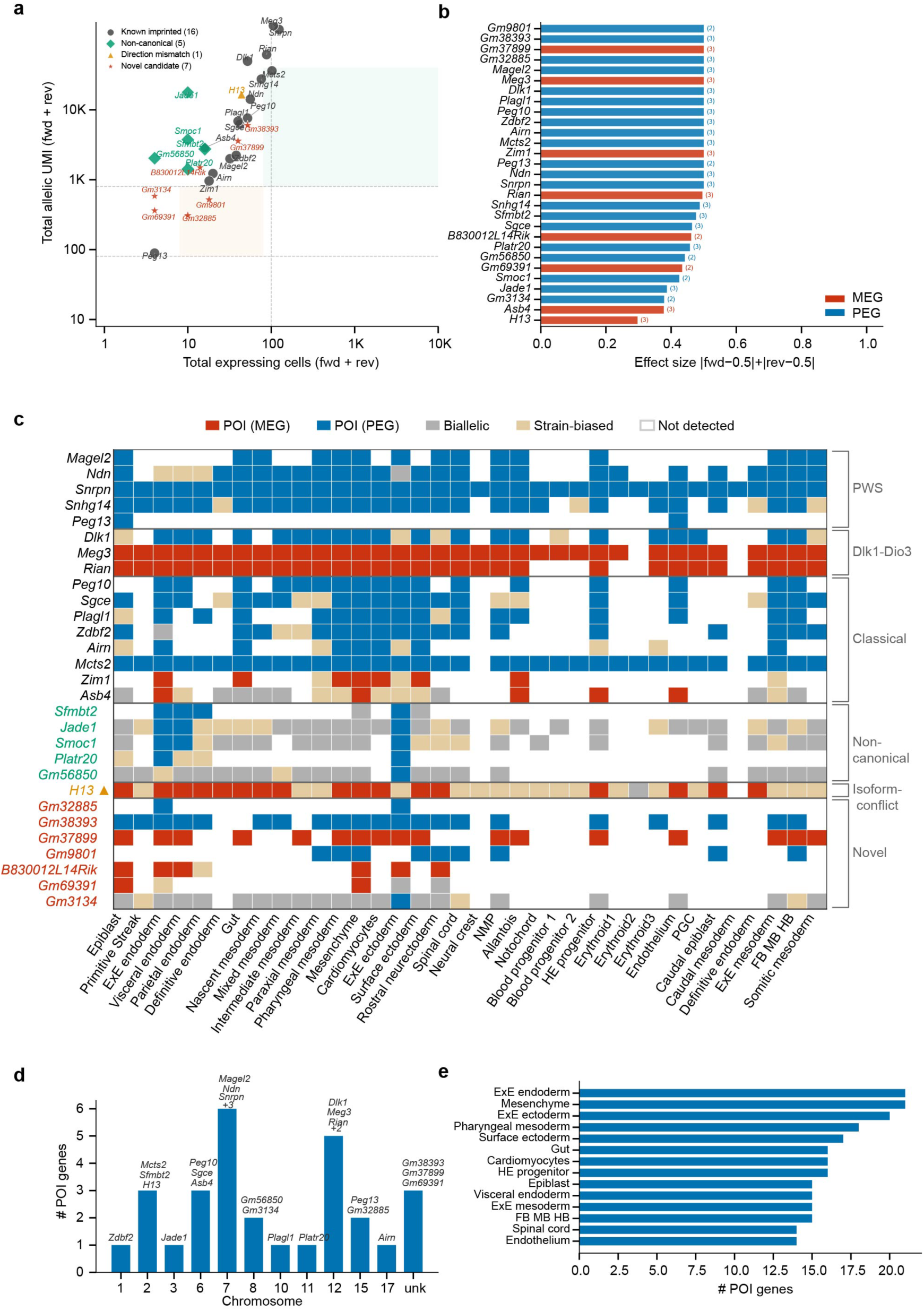
| Quantitative evidence and developmental distribution of the 29-gene CT-POI panel. (**a**) Per-gene expressing cells versus allele-informative UMIs (both crosses, log scale; shape, category; area, effect size; bands, evidence strength). (**b**) Genes ranked by effect size (|forward AR − 0.5| + |reverse AR − 0.5|); colour, direction; parenthetical numbers, stages with reciprocal flip. (**c**) Gene × cell-type matrix (POI-maternal, POI-paternal, biallelic, strain-biased, not detected). (**d**) CT-POI genes per chromosome. (**e**) Cell types ranked by number of CT-POI genes (top 14).

**Supplementary Fig. 2.**
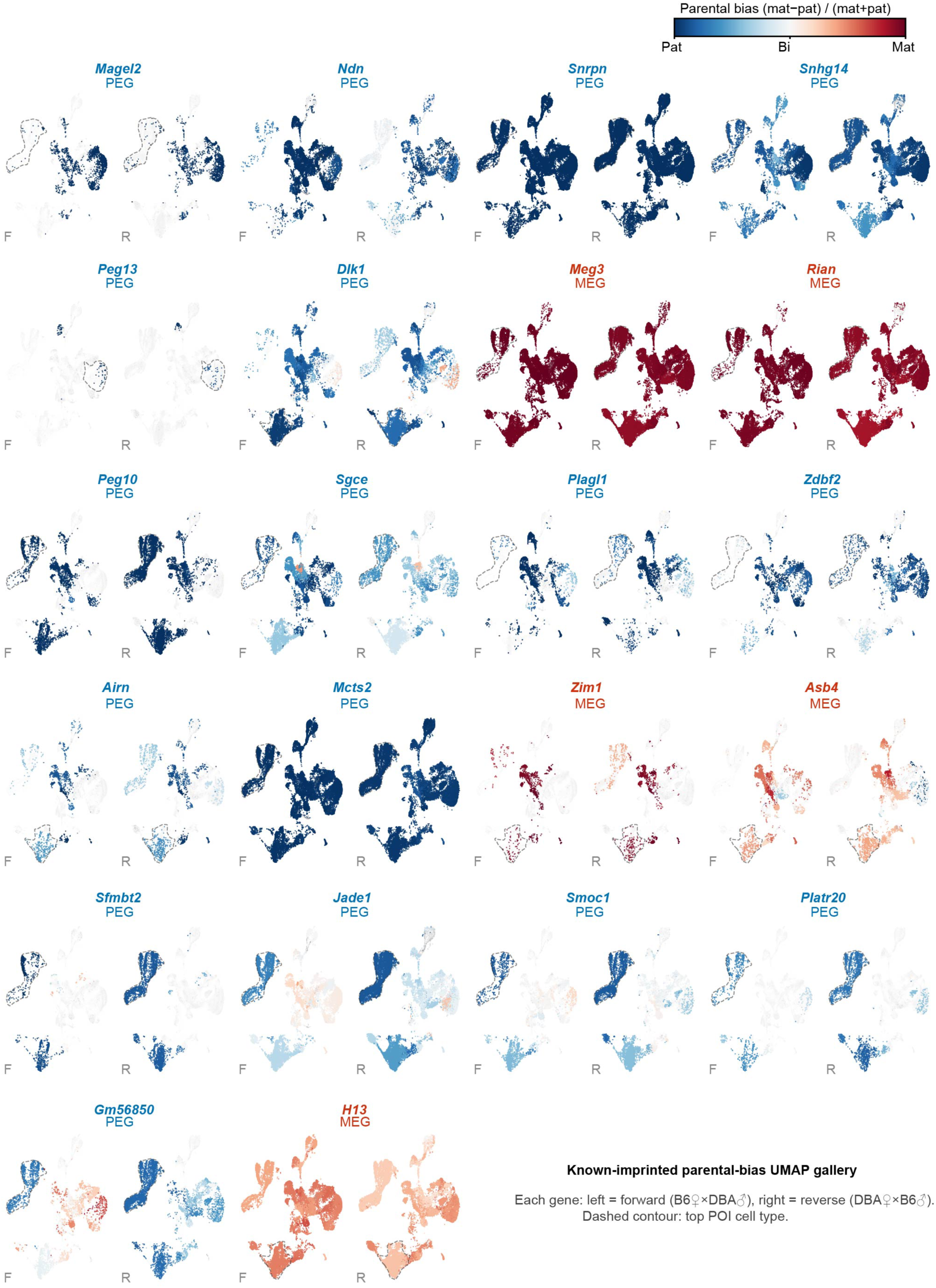
| Reciprocal-cross parental-bias UMAP gallery for 22 known imprinted genes. Paired forward and reverse maps per gene, coloured by per-cell-type parental-bias score; grey, insufficient informative UMIs; dashed contour, strongest-signal cell type. A genuine parent-of-origin effect appears as a colour inversion between crosses; a strain-fixed effect retains the same colour. Genes ordered by imprinted locus.

**Supplementary Fig. 3.**
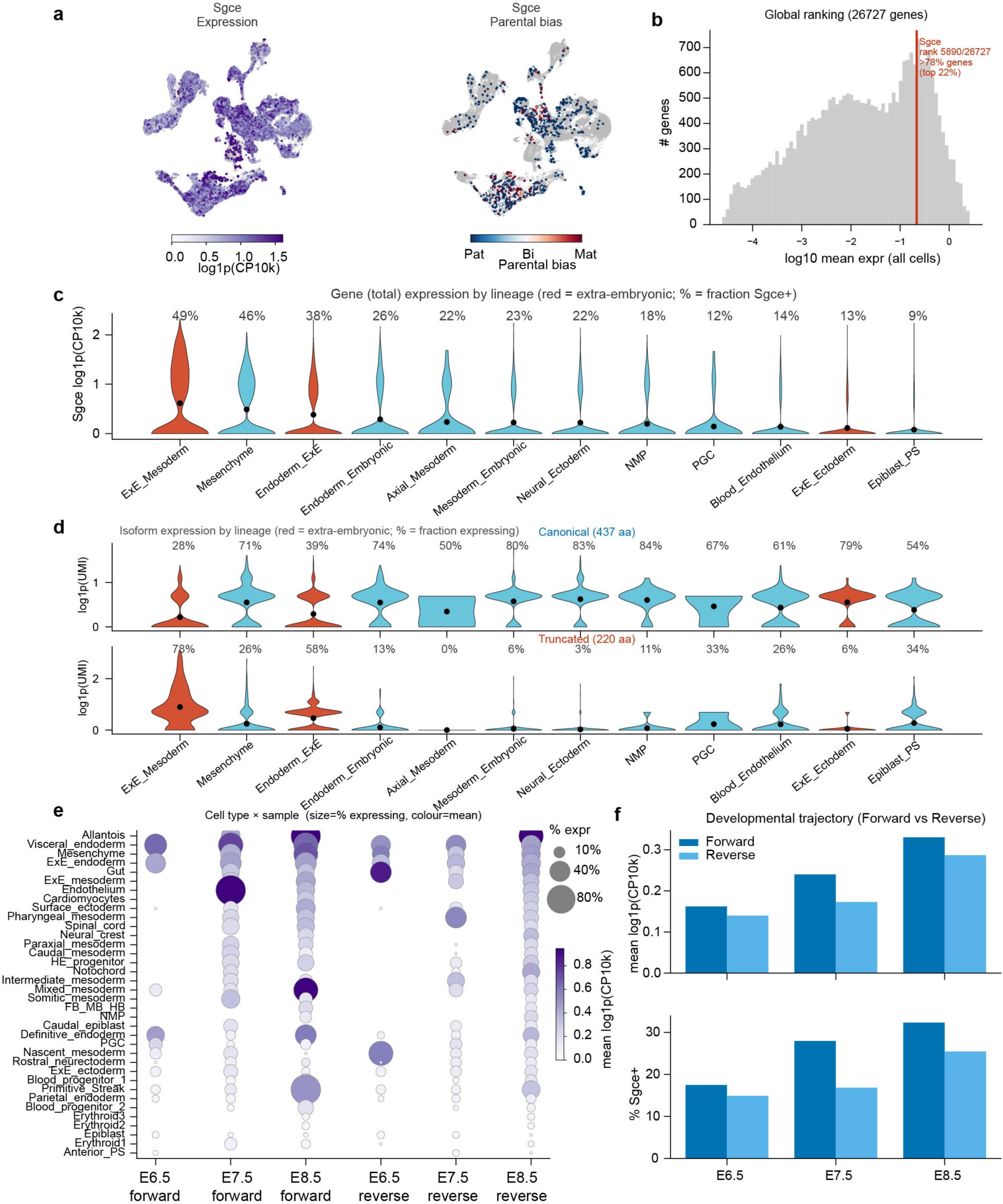
| *Sgce* expression across gastrulation at the gene and isoform level. (**a**) UMAP coloured by *Sgce* total expression and gene-level parental bias. (**b**) Distribution of mean expression across genes (red line, *Sgce*). (**c**) *Sgce* expression across lineages (extra-embryonic in red; % expressing above each violin). (**d**) Canonical (437-aa) versus truncated, L1MC1-derived (220-aa) isoform expression across lineages. (**e**) Dot plot per fine cell type and sample. (**f**) Mean expression and detection rate per stage, forward versus reverse.

**Supplementary Fig. 4.**
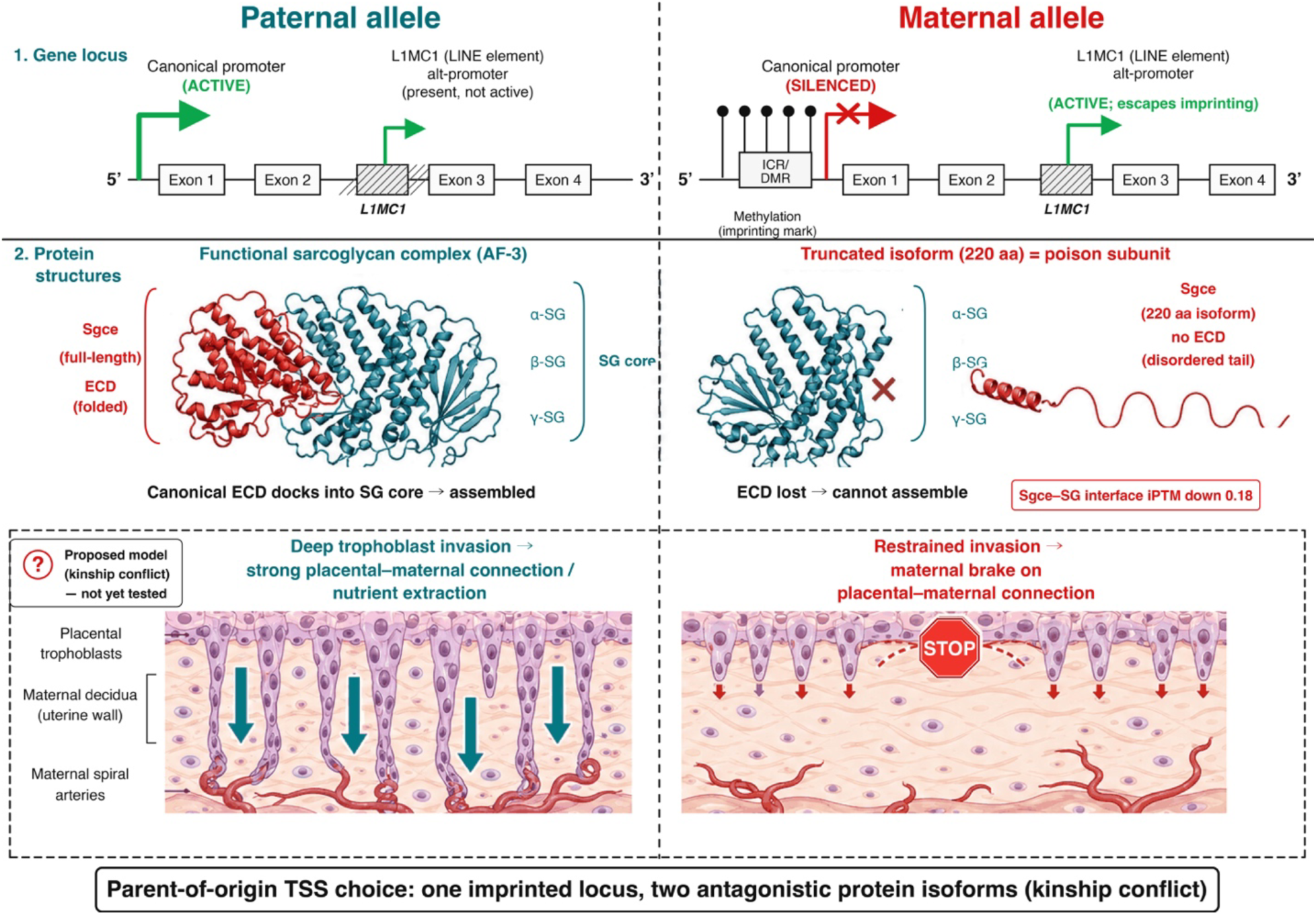
| Proposed kinship-conflict model for *Sgce* L1MC1 imprinting (hypothesis). Schematic of parent-of-origin transcription-start-site choice: the paternal canonical promoter yields an assembling sarcoglycan complex, whereas the maternal L1MC1 alternative promoter yields a truncated, non-assembling “poison” subunit; a proposed maternal brake on trophoblast invasion is marked explicitly as an untested hypothesis.

**Supplementary Tables 1–7** as listed in the submission inventory.

**Supplementary Note 1 — ANCHOR algorithmic specification**

*This note gives the implementation detail required to reimplement ANCHOR. Symbols match the main text: molecule-by-site sign matrix A ∈ {+1, −1, NaN}, molecule haplotype vector h, site phase vector δ*.

**1.** **Signed-graph rank-1 factorization (de-novo caller front end).** For each chromosome, A is built from per-UMI consensus reads: entry A_{m,s} = +1 if molecule m matches the reference base at site s, −1 if it matches the alternate base, NaN if uncovered. We minimise the weighted reconstruction objective J(h,δ) = Σ_{(m,s): observed} w_{m,s} (A_{m,s} − h_m δ_s)² over h_m, δ_s ∈ {−1,+1}, where w_{m,s} is the base-quality-derived confidence (w = 1 − 10^(−Q/10)); missing entries are excluded from the sum (no imputation). h is initialized from the sign of the leading left singular vector of the observed-masked, weight-scaled A (spectral initialization). Coordinate updates alternate to the weighted-majority sign: h_m ← sign(Σ_s w_{m,s} A_{m,s} δ_s) and δ_s ← sign(Σ_m w_{m,s} A_{m,s} h_m), iterated to convergence (flip rate < 10^−4, typically ≤ 8 iterations). Disconnected components of the molecule–site graph are factorized independently and assigned arbitrary but internally consistent global sign; phase-set boundaries are placed at component boundaries and at sites with no linking molecule. Per-site features derived from the factorization include frustration f_s = Σ_m w_{m,s} 1[A_{m,s} ≠ h_m δ_s] / Σ_m w_{m,s} (the weighted fraction of a site’s observations that disagree with the rank-1 reconstruction h_m δ_s) and graph negative mass g_s = Σ_m w_{m,s} 1[A_{m,s} ≠ h_m δ_s] (the same weighted disagreement, un-normalized). Complexity is O(nnz) per iteration (nnz = non-missing entries); memory uses a CSR sparse representation, processed per chromosome. Indels are represented as a collapsed multi-base allele at the left-aligned position; sites that cannot be left-aligned uniquely are excluded.
**2.** **Classifier cascade.** Stage 1 (BAM/context) and stage 2 (graph-aware reranker) are LightGBM models on 34 features (Supplementary Table S4: read-level BAM statistics, derived quality/concordance, single-cell support counts, graph/linkage features, sequence/RNA-editing context, haplotype Bayes factors). Hyperparameters: num_leaves = 63, max_depth = 8, learning_rate = 0.03, n_estimators = 800, no early stopping, no scale_pos_weight (probabilities span [0,1] for calibrated operating points). Training used chromosome 1–10 for HG002 and leave-one-chromosome-out for mouse; the test chromosome is never seen in training or validation (no site leaks across the split). Production operating point s2 ≥ 0.70 (s2 = the stage-2 classifier probability); high-precision mode s2 ≥ 0.95. Calibration (reliability, Extended Data Fig. 2a) is computed per held-out chromosome and reported pooled; the model probability is used directly (no post-hoc recalibration), and its conservativeness is disclosed.
**3.** **Pair-HMM emission and read-level HMM (allelic engine).** The pair-HMM aligns each read to each of the two haplotype reference sequences over states {M (match/mismatch), I_read (read insertion), I_ref (reference insertion/deletion)} with affine gap transitions; emission at M uses the per-base error probability from the ONT base quality (e = 10^(−Q/10)), with a homopolymer-length-dependent inflation term for runs ≥ 5 nt; splice gaps (N-CIGAR) are treated as zero-cost reference jumps, not insertions. The per-site LLR is the difference of forward log-likelihoods under the two haplotypes restricted to the window around each informative site. LLRs are calibrated by Platt logistic regression with batch-aware covariates (sample, run); calibration is fit per sample and shared across chromosomes. The read-level HMM integrates calibrated per-site LLRs along the molecule over states {hap1, hap2, switch} with a fixed switch penalty; a UMI’s reads are aggregated by a Beta-binomial (overdispersion ρ = 0.01) into a four-state posterior {hap1, hap2, MIX, UNK}, abstaining (MIX/UNK) when neither haplotype posterior exceeds 0.95.
**4.** **Isoform assignment.** Reads are grouped into splice chains by ordered junction coordinates with a ±10-bp tolerance at each junction; a chain requires ≥ 2 supporting reads to define an isoform. Transcript 5′/3′ ends are not used to distinguish isoforms (10x cDNA does not define them); UMIs are collapsed to one isoform call by majority of their reads’ chains. Isoforms are equivalence classes of junction chains; allele-specific junctions are retained as distinct isoforms even when they map to the same reference transcript. Reference annotation (GENCODE/Ensembl) is used only to *label* isoforms, never to *seed* or constrain chain discovery, so allele-specific splicing is preserved.
**5.** **Benchmark operating definitions.** “Callable truth” at minimum coverage N = GIAB SNPs ∩ (BAM depth ≥ N, MAPQ ≥ 5, BQ ≥ 10), independent of any caller. Competitors were run PASS-only on the identical RNA BAM with no genotype or phased VCF input; for het/ASE comparisons their calls were restricted to heterozygous SNPs within confident regions. Exact versions, commands, filters and output fields are in Supplementary Table S5.
**6.** **Numerical constants and software (for exact reimplementation).** Pair-HMM: affine gaps with gap-open = 10^−3 and gap-extend = 0.4 per position; match-state emission uses the per-base error e = 10^(−Q/10) from the ONT quality string, with homopolymer inflation e’ = min(0.5, e·(1 + 0.25·max(0, L−4))) for a run of length L; splice gaps (CIGAR N) are zero-cost reference jumps bounded to annotated or ≥ 20-bp gaps; the per-site LLR is taken over a ±25-bp window around each informative site. Platt calibration: logistic regression of the correct haplotype label on the raw per-site LLR plus indicator covariates for sample and sequencing run; fit per sample, shared across chromosomes. Read-level HMM: states {hap1, hap2, switch}, switch penalty = 10^−4 per junction, posterior decision threshold 0.95; UMI Beta-binomial overdispersion ρ = 0.01; minimum 1 informative read per UMI and ≥ 1 informative UMI per (cell, gene) for a call. Software: LightGBM 4.3.0, Python 3.10, fixed random_state = 0 with deterministic = true and force_row_wise = true; positive class = GIAB/marker-panel true heterozygous site, negative class = non-truth candidate; model selection by validation AUPRC; features are encoded identically across human, mouse and bulk runs (no per-dataset re-encoding). Isoform-assignment edge cases: ambiguous splice chains (a read compatible with > 1 chain) are assigned to the highest-support chain and dropped if the top two are within 10%; partial 3′-biased molecules contribute only to chains whose covered junctions they fully support; single-exon transcripts are assigned by overlap to a single equivalence class; retained introns are treated as distinct isoforms when the intron is spanned by ≥ 2 reads; junctions with < 2 supporting reads are not used to define isoforms; reads conflicting with the UMI-majority chain are excluded from that UMI’s isoform call.

